# Sonoepigenetic Modification Mechanoprimes Early Osteogenic Commitment in Mesenchymal Stem Cells

**DOI:** 10.1101/2025.03.17.643773

**Authors:** Lizebona A. Ambattu, Blanca del Rosal, Carmelo Ferrai, Leslie Y. Yeo

**Affiliations:** Micro/Nanophysics Research Laboratory, School of Engineering, RMIT University, Melbourne, VIC 3001, Australia; School of Science, RMIT University, Melbourne, VIC 3001, Australia; Institute of Pathology, University Medical Center Göttingen, Robert-Koch-Str. 40, 37075, Göttingen, Niedersachsen, Germany

**Keywords:** nuclear mechanotransduction, calcium signaling, cAMP, RNA polymerase II, histone modification

## Abstract

Cells effectively balance and integrate numerous pathways to adapt to external signals in an attempt to regain homeostasis, although the complex nuclear mechanotransduction mechanism through which this occurs is not as yet fully understood. Contrary to prevalent thought that the relay of extracellular cues to the nucleus to effect its fate and function predominantly relies on direct transmission through the cytoskeletal structure, we demonstrate, through the use of high frequency (10 MHz) nanomechanostimulation, that induced fluctuations of the cells’ nuclear chromatin response are primarily influenced by the spatiotemporal dynamics associated with the bidirectional crosstalk between two key second messengers, namely calcium (Ca^2+^) and cyclic adenosine monophosphate (cAMP). We show that this conditioning is an adaptive response to the mechanostimuli and correlates with a ‘mechanopriming’ effect. Notably, brief (10 mins) daily exposure to the mechanostimulation was sufficient to direct mesenchymal stem cells toward an osteogenic lineage in as little as three days—without the need for osteogenic factors.

## Introduction

There is growing recognition that the interplay between physical cues and their impact on cell homeostasis, mediated by nuclear mechanics and chromatin regulation, are central to virtually all physiological and developmental processes [1, 2, 3]. It has been observed, for example, that mechanical perturbations can induce changes in the distribution of Lamin A/C at the nuclear periphery and wrinkling of the nuclear membrane, implying the mechanotransductive role of the cytoskeletal structure in relaying signals from the extracellular environment to the nuclear membrane through the LINC (linker of nucleoskeleton and cytoskeleton) complex [4, 5, 6, 7]. Additionally, mechanical forces have also been found to drive chromatin changes by reorganizing the epigenomic landscape through histone modifications [3, 8, 9], enabling cells to finely tune gene expression in response to external cues to maintain homeostasis [10, 11]. The specific molecular mechanisms by which extracellular cues from the cells’ periphery are transmitted to the nucleus to influence chromatin organization, and how these aspects affects cell fate, however, remain poorly understood [3].

The discrepancy in the timescales associated with the nuclear mechanical responses—with nuclear deformation occurring over millisecond timescales [12, 13], and chromatin reorganization taking seconds to minutes [14, 15, 16]—nevertheless hints at a more complex mechanotransduction picture. The involvement of Ca^2+^-responsive kinases and phosphatases that influence numerous epigenomic and chromatin regulators, for instance, suggests that other, and possibly multiple, mechanisms could be involved in the regulation of chromatin and epigenomic landscapes in response to the extracellular cues [17, 18, 19, 20, 21, 22]. That Ca^2+^-induced signaling pathways activated by external stressors to the cell are typically counterbalanced by other signaling cascades involving another second messenger cyclic adenosine monophosphate (cAMP) during the restoration of the cell to its homeostatic state, suggests that cAMP may also play a pivotal role in the dynamics of epigenetic regulation associated with the recovery process [23, 24].

The way by which transmission of mechanical forces through the cytoskeleton dynamically interplays with these biochemical signals (such as Ca^2+^[20, 21, 25, 26, 27, 28, 29] and cAMP [23, 24]) over different time scales to facilitate latent recovery of the cell towards homeostasis (and to ultimately influence transcriptional outcomes and cell fates [30], including those related to stem cell differentiation and function [31, 32, 33]) remains to be explicated. To this end, we have found it expedient to utilize high frequency (10 MHz) nanomechanostimulation in the form of nanometer-amplitude hybrid surface and bulk sound waves (surface reflected bulk waves (SRBW) [34]) to examine the nuclear mechanotransduction process. Besides the physiological relevance of this specific frequency for activating mechanosensitive ion channels [35, 36], the low amplitude high frequency mechanostimulation has been shown to facilitate spatiotemporal modulation of Ca^2+^ and cAMP within the cell reproducibly with a fine degree of control that does not drive the cell toward apoptosis [37]; such an approach therefore differing from other conventional forms of ultrasonic stimulation, particularly at lower kHz order frequencies, where cavitation, heating and acoustic streaming can typically limit the ability to modulate Ca^2+^ into the cell and hence its recovery towards homeostasis [38]. In particular, we show that such spatiotemporal Ca^2+^ dynamics initiates a biphasic nuclear response in the cells associated with the initial challenge and subsequently their recovery, thereby offering a window by which insight can be gleaned into the multi-timescale role and bidirectional crosstalk between Ca^2+^ and cAMP in the dynamics of nuclear and epigenetic regulation.

More specifically, we unravel the role of the cytoskeletal structure and second messenger signaling in the nuclear mechanotransduction process. Tension triggered by the actin cytoskeletal rearrangement in response to the SRBW nanomechanostimulation is observed to drive transient modifications in nuclear morphology and Lamin dynamics. Interestingly, we find that the mechanostimulation also induces a fluctuation on several histone modifications. Functional experiments with specific inhibitors show that such a response is specifically controlled by the dynamic interplay between Ca^2+^ and cAMP signaling, independent of actin cytoskeletal activity. Further, we note that, together with the induced fluctuation of H3K27me3, H3K9me3 and H3K27ac, the mechanostimuli promoted clustering of RNA polymerase II (RNAPII) and Cajal bodies (CBs), thereby revealing for the first time, the mechanoresponsive behavior of these nuclear bodies as a result of the second messenger dynamics.

Altogether, we elucidate how the SRBW-induced epigenetic regulation dynamics are influenced by spatiotemporal changes in intracellular Ca^2+^ and cAMP levels. In particular, we observe the Ca^2+^–cAMP signaling to be intricately related to the process by which mesenchymal stem cells are transcriptionally primed for early commitment towards osteogenic differentiation [39]. Importantly, inhibition of such second messenger dynamics was observed to suppress early osteogenic lineage induction of the cells, demonstrating that the SRBW-induced regulation of chromatin fluctuations is not merely causal, but occurs downstream of the Ca^2+^–cAMP signaling cascade.

## Results and Discussion

### Nuclear Mechanoresponse

Subjecting bone-marrow-derived human mesenchymal stem cells (hMSCs) to 10 mins of 2.5 W SRBW nanomechanostimulation (optimized to achieve the observed effect in the cells whilst maximizing their viability (≈ 95%)), as shown in the experimental setup described in the Methods section and depicted in Fig. 1a and Fig. S1 (Supplementary Information), can be seen to induce a biphasic response (Fig. 1b). First, we observe an immediate surge in Ca^2+^ within the cell by approximately 2.3 times (Fig. 1b,c), enabled by activation of mechanosensitive Piezo1 channels as a consequence of the mechanostimulation (i.e., the *sonochallenge* phase) [40]. Following this spike, the intracellular Ca^2+^ levels returned to their baseline values within 10 mins of post-exposure incubation (the 0 min post-exposure time point corresponding to the instant at which cells were fixed immediately after the 10 min nanomechanostimulation duration). As a secondary latent response to the mechanostimulation, we also observed a steady increase in intracellular cAMP levels at around 15 mins post-exposure, reaching a peak within an hour, at which they remained elevated before gradually returning to baseline levels after 8 hrs (i.e., the *sonotransformation* phase; Fig. 1b,d). Parenthetically, we note that although G protein–coupled receptors (GPCRs) are well-established upstream regulators of the cAMP pathway and are known to be responsive to membrane tension, our findings indicate that the subsequent cAMP response observed here is primarily mediated through Piezo1-activated Ca^2+^ influx rather than GPCR signaling [37, 39, 40]).

**Figure 1:**
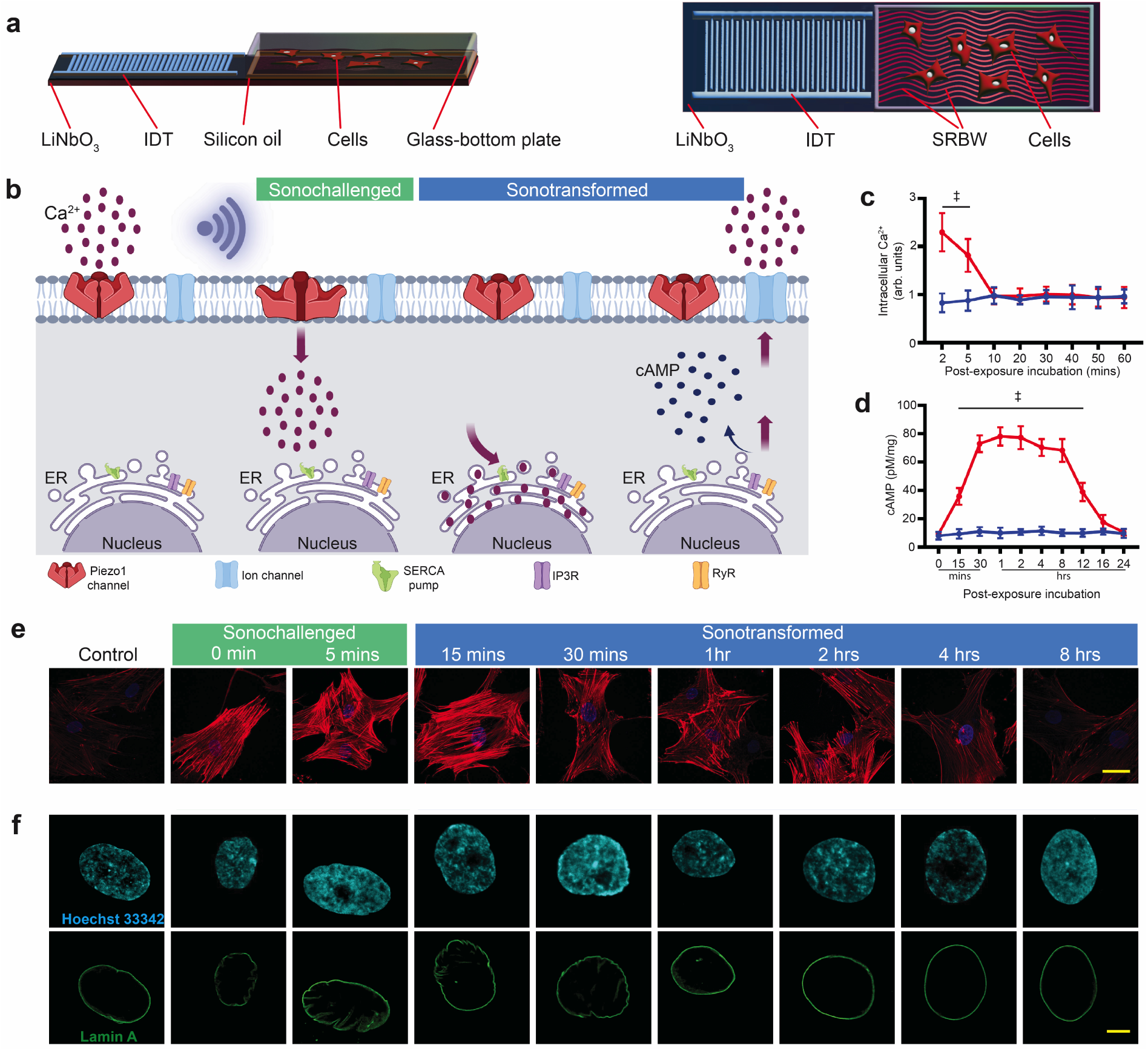
SRBW nanomechanostimulation induces nuclear deformation and Lamin A wrinkling in hMSCs. (a) Side (left) and top (right) view schematic representations of the experimental setup in which the SRBW (not to scale; see also image of the setup in Fig. S1 (Supporting Information)), produced by applying a sinusoidal electrical signal at its resonant frequency (10 MHz) to an interdigitated transducer (IDT) photolithographically patterned on a single crystal piezoelectric substrate (LiNbO_3_), is transmitted through a thin layer of silicon oil into a glass-bottom culture plate containing adherent bone-marrow-derived mesenchymal stem cells (hMSCs). (b) Schematic illustrating the Ca^2+^ mobilization pathways within the cell throughout the time frame associated with the biphasic response of the cell to the SRBW nanomechanostimulation (see Fig. S2 (Supporting Information)): from left to right: control or pre-stimulation condition, the sonochallenge phase following stimulation, and the sonotransformation phase associated with the recovery stage of the cell towards homeostasis. (c) Intracellular Ca^2+^ (*n* = 4) and (d) cAMP (*n* = 4) concentration in the hMSCs as a function of post-exposure incubation time for both control (unstimulated; blue datapoints) and SRBW nanomechanostimulated (red datapoints) cells; data are represented in terms of the mean value ± the standard error over multiple runs, and, ‡ indicates statistically significant differences with *p <* 0.0001. (e,f) Representative confocal microscopy images illustrating (e) the reorganization of the actin cytoskeleton (60× magnification; scale bar: 50 µm), and, (f) the nucleus (100× magnification; scale bar: 5 µm) in hMSCs subjected to the SRBW nanomechanostimulation at different post-exposure incubation times compared to the control (unstimulated) group. Cell nuclei were stained with Hoechst 33342 (displayed in cyan) and Lamin A immunofluorescence (displayed in green), whereas F-actin was labeled using ActinRed™ 555 (red).

Over this time course, a rapid, pronounced increase in actin stress fibers over the nuclear region is apparent (Fig. 1e and Fig. S3 (Supporting Information)). This characteristic cap can be seen to persist up to 15 mins following the SRBW nanomechanostimulation, commensurate with the timescale of the initial Ca^2+^ spike transient associated with the sonochallenge. At 30 mins, the actin organization is already focused toward the periphery of the cell, and after 8 hrs, the cell gradually returns to its homeostatic state, similar to the unstimulated control (see Fig. 1e). Morphometric measurements of the nucleus with Lamin A staining (Fig. 1f) showed that while no appreciable changes were observed in the total nuclear volume (Fig. S4a (Supporting Information)), the SRBW nanomechanostimulation nevertheless induced significant alterations to the nucleus’ general shape. This is highlighted by the decrease in its transverse area (Fig. S4b (Supporting Information)) and an increase in its height (Fig. S4c (Supporting Information)), with consequent changes in its circularity, solidity and eccentricity (Fig. S4d,e,f (Supporting Information)). Additionally, we observe Lamin A to transition from its initially smooth, oval shape to an undulated and wrinkled appearance (Fig. 1f).

The time-course analysis of the images indicates the alignment of the observed evolution in morphological transformation with the biphasic Ca^2+^ and cAMP time scales in Fig. 1c,d. Rapid and sharp morphological changes over the timescale (minutes) over which Ca^2+^ levels are first elevated following the initial sonochallenge phase—characterized by stress fiber formation and reduction in nuclear size. This was then followed by their gradual recovery (size, area, volume, and shape descriptors) over the timescale (hours) associated with the subsequent sonotransformation phase during which the cell relaxes back to its ground state (Fig. S2 (Supporting Information)).

Alterations in nuclear morphology, particularly those involving nuclear Lamins, have been reported to affect chromatin organization, leading to modifications in chromatin condensation and accessibility [6, 8, 10, 41, 42]. Considering the effects of the SRBW nanomechanostimulation on the cells, we assessed the distribution of H3K27me3 (Fig. 2a), a facultative heterochro-matin marker mediated by EZH2 (a component of the Polycomb repressive complex PRC2), and the constitutive heterochromatin marker H3K9me3 (Fig. 2b). The results show an initial slight downregulation of H3K27me3, which precede significant upregulation upon normalization of cytosolic Ca^2+^ levels and elevation of cAMP during the sonotransformation phase at 30 mins. More specifically, H3K27me3 increases by approximately fivefold before gradually decreasing back to baseline levels with the recovery of the cell over 4–8 hrs (Fig. 2c, and Figs. S5 and S6c (Supporting Information)). Interestingly, the immunofluorescence for H3K9me3 shows a different dynamic and is somehow anticorrelated with the H3K27me3 nuclear distribution, exhibiting a decrease to approximately 0.2-fold at 30 mins before gradually regaining its signal to baseline levels with the recovery of the cell over 4–8 hrs (Fig. 2d). Examination of a chromatin marker for active enhancers H3K27ac [43] in Fig. 3a, and Figs. S6a,b and S7 (Supporting Information), on the other hand, showed its significant upregulation upon normalization of cytosolic Ca^2+^ levels and elevation of cAMP during the sonotransformation phase at 30 mins by approximately threefold, followed by a return to baseline levels over 4–8 hrs (Fig. 3d, and Figs. S6a,b and S7 (Supporting Information)).

**Figure 2:**
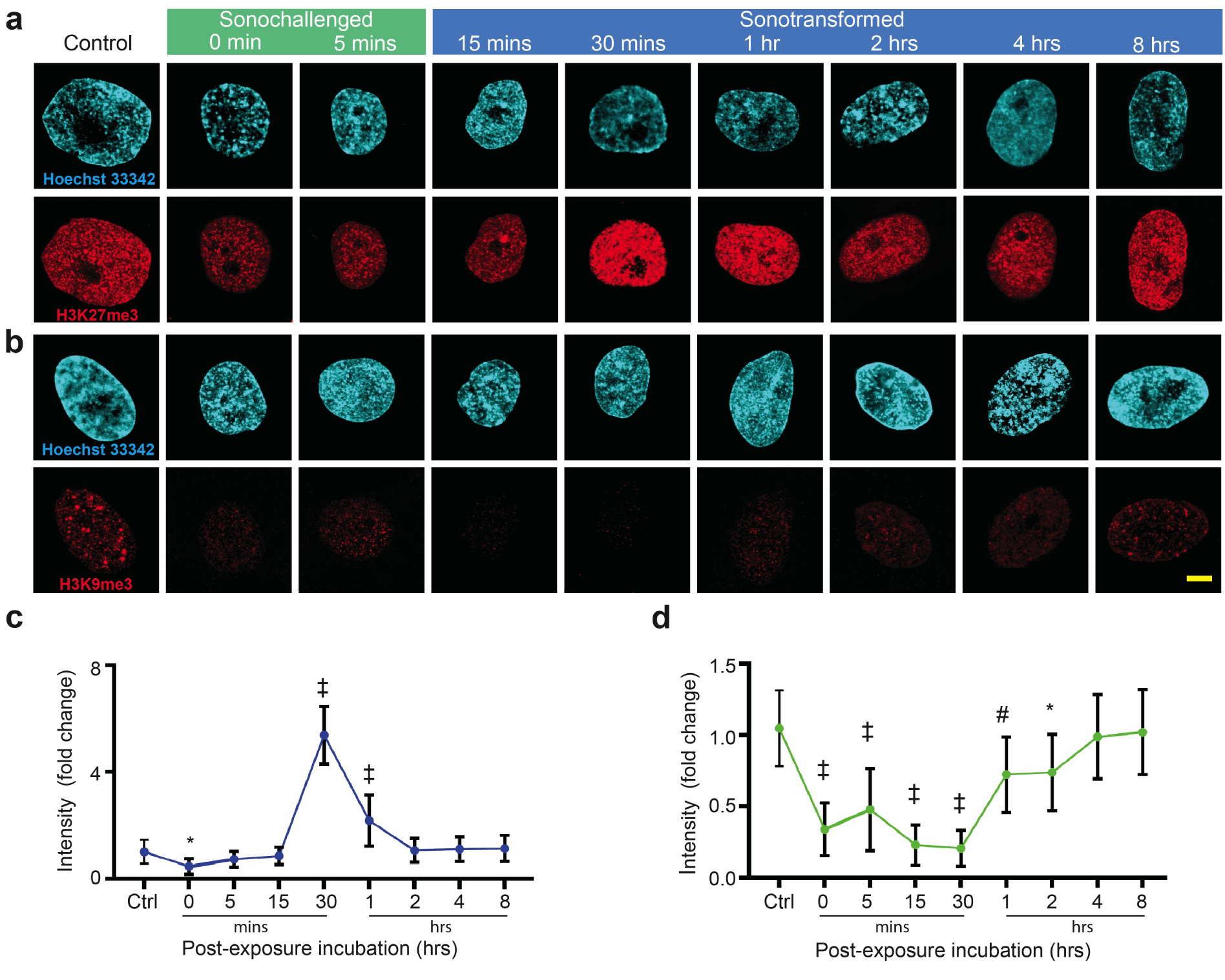
hMSCs show fluctuations in chromatin repressive marks in response to the SRBW nanomechanostimulation. Representative confocal microscopy images of the nucleus (stained with Hoechst 33342 and displayed in cyan) together with (a) H3K27me3 and (b) H3K9me3 (displayed in red) immunofluorescence following SRBW nanomechanostimulation of the hMSCs at different post-exposure incubation times, compared to those of the control (unstimulated) cells. All images were acquired at 100× magnification and the scale bars denote lengths of 5 µm; lower magnification images showing similar representative observations in a larger number of cells can be found in Fig. S5 (Supporting Information). Also shown are the relative immunofluorescence of (c) H3K27me3 and (d) H3K9me3 in the SRBW nanomechanostimulated cells at different post-exposure incubation times (0 min corresponding to the time point at which cells were fixed immediately after the 10 min nanomechanostimulation duration) with respect to that in the control (unstimulated) cells. Data are represented in terms of the mean value ± the standard error over multiple runs (*n >* 50 cells/condition, visualized across four independent experiments); ∗, # and ‡ indicate statistically significant differences with *p <* 0.05, *p <* 0.01 and *p <* 0.0001, respectively.

**Figure 3:**
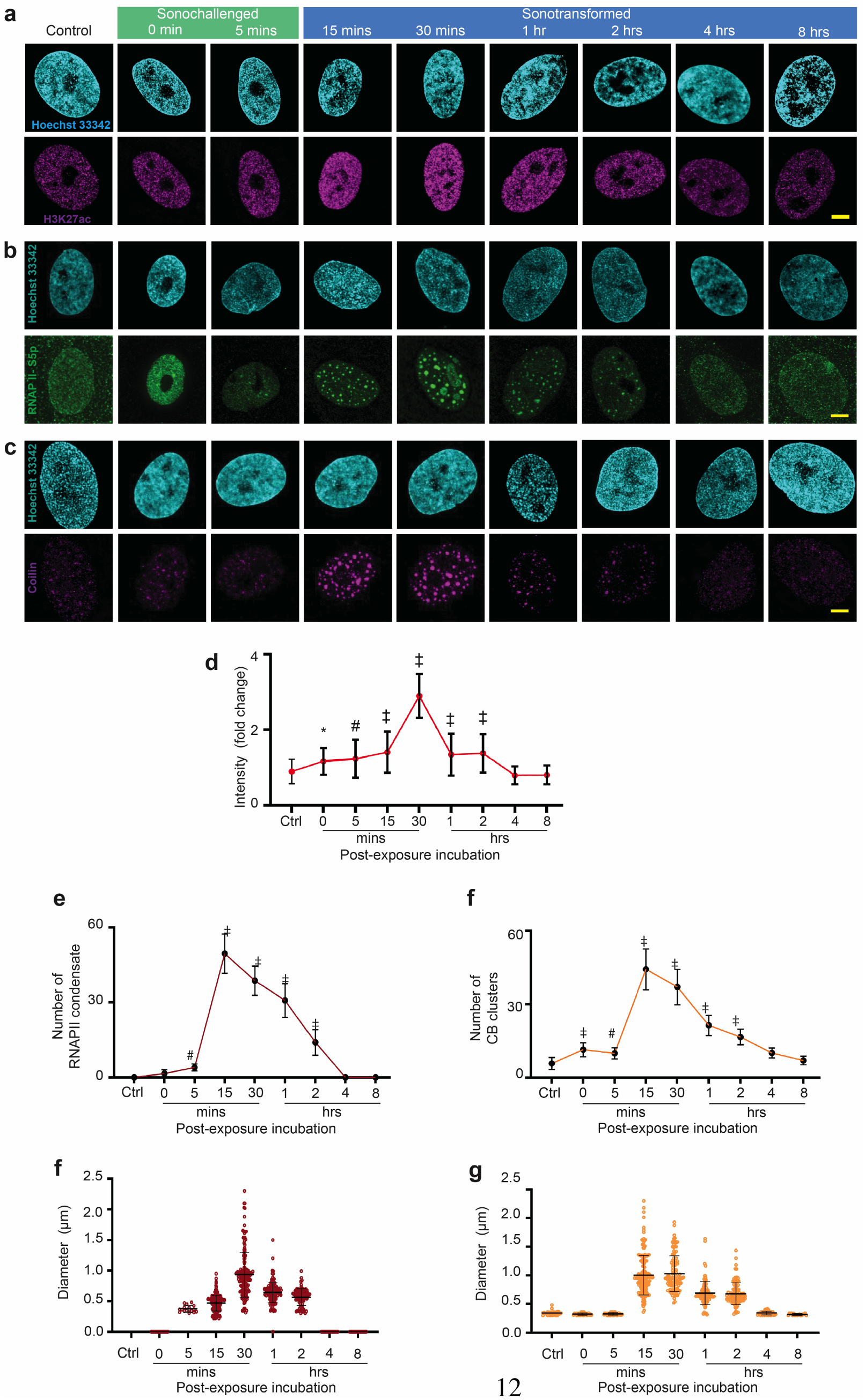
Nuclear bodies exhibit mechanoresponsive behavior to SRBW nanomechanostimulation. (a–c) Representative confocal microscopy images of of the nucleus (stained with Hoechst 33342 and displayed in cyan) together with (a) H3K27ac (displayed in purple), (b) RNAPII-S5p (displayed in green), and, (c) Coilin (displayed in purple) immunofluorescence following SRBW nanomechanostimulation of the hMSCs at different post-exposure incubation times, compared to those of the control (unstimulated) cells. All images were acquired at 100× magnification and the scale bars denote lengths of 5 µm. Also shown are (d) the relative immunofluorescence of H3K27ac, (e) the number, and, (f) the size of the RNAPII-S5p condensates, respectively, and, those of (g,h) the Coilin clusters, respectively, in the SRBW nanomechanostimulated cells at different post-exposure incubation times. Data are represented in terms of the mean value ± the standard error over multiple runs (*n >* 50 cells/condition, visualized across four independent experiments); ∗, # and ‡ indicate statistically significant differences with *p <* 0.05, *p <* 0.01 and *p <* 0.0001, respectively.

Strikingly, staining for RNAPII CTD Serin5 phosphorylation (RNAPII-S5p)—a specific post-translational modification that characterizes transcriptionally engaged RNAPII (active and poised) [44, 45]—revealed a dramatic pattern distribution accompanying the spike in intracellular Ca^2+^ during the initial sonochallenge phase (Fig. 3b, and Fig. S6d (Supporting Information)). In contrast to active RNAPII molecules which are typically localized within distinct but adjacently-placed spatial compartments with approximate lengthscales on the order 100 nm [46, 47], the SRBW nanomechanostimulation can instead be observed to induce an increase in the number of the RNAPII-S5p larger clusters by 15 mins (Fig. 3e), which progressively grew into bigger condensates (Fig. 3f). We ruled out that these condensates could have resulted from a block in transcription as a consequence of DNA damage induced by the SRBW nanomechanostimulation, as demonstrated by the absence of histone H2AX phosphorylation—a marker for early cellular responses to DNA damage [48]—in the treated cells (Fig. S8 (Supporting Information)).

Similar to RNAPII-S5p staining, we also observed the immunofluorescence signal for Coilin—a marker of Cajal Bodies (CBs)—to also transiently cluster towards bigger condensates by 15 mins (Fig. 3c,h and Fig. S7 (Supporting Information)), suggesting the broader effect of the SRBW nanomechanostimulation on inducing alterations to other nuclear condensates. Quantitative analysis of the CB clusters shows that they increased in their dimension from 0.3 µm at 5 mins before reaching a peak of 1.1 µm at 30 mins (Fig. 3h), albeit with decreasing number of condensates (Fig. 3g). Quite interestingly, the results reveal a lack of coincidence between the total numbers and dimensions of RNAPII-S5p and the CBs, suggesting that the two aggregates could represent independent functions despite a certain level of interaction. Akin to the other chromatin features, it can also be seen that the RNAPII-S5p and CB expression gradually returned to their original size upon recovery of the cell during the sonotransformation phase over which intracellular cAMP levels were elevated.

### Role of Spatiotemporal Ca^2+^ and cAMP Dynamics

The aforementioned results highlight the biphasic response of the cells toward the SRBW nanomechanostimulation, through which the spatiotemporal dynamic interplay between Ca^2+^ and cAMP in modulating nuclear morphology and the chromatin fluctuations becomes evident. (1) An initial phase where the Piezo1-mediated influx of Ca^2+^ into the cell, upon being *sonochallenged* by the SRBW nanomechanostimulation, triggers various signaling pathways involving the small guanosine triphosphate hydrolase (GTPase) Rho (Ras homolog gene family) and its downstream effector ROCK (Rho-associated protein kinase), which are responsible for the formation of the stress fibers observed in Fig. 1e and Fig. S3 (Supporting Information) [49, 50, 51] up to around 15 mins post-exposure. (2) A subsequent *sonotransformation* phase after approximately 30 mins in which the stress fibers are dissolved upon activation of the cAMP-activated guanine nucleotide exchange protein/Ras-related protein 1 (Epac/Rap1) signaling cascade driven by the elevated cAMP concentration [37, 52, 53, 54, 55]. This biphasic behavior (Fig. 1b), which is a typical with the SRBW nanomechanostimulation [37], starkly contrasts with the classical monophasic response of Ca^2+^-induced stress fiber formation following challenges to the cell either with chemical or conventional (low frequency (<1 MHz)) mechanical stimulation [56, 57, 58]. This contrasting behavior is likely due to the distinctive capacity of the SRBW to facilitate bidirectional crosstalk between the spatiotemporal Ca^2+^ dynamics and opposing cAMP signaling, given its ability to modulate sufficient levels of Ca^2+^ into the cell without driving the cells toward apoptosis. This allows excess Ca^2+^ to be sequestered into the endoplasmic reticulum (ER) such that its subsequent latent release back into the cytosol through ryanodine (RyR) channels and inositol trisphosphate (IP3R) receptors (calcium-induced calcium release; CICR [59]) over considerably longer time periods (typically hours) enables cAMP signaling and hence Epac/Rap1 counteraction as part of the recovery process by which the cells return to their homeostatic state [37].

Specific inhibitors were adopted to glean further insight into the mechanotransduction mechanism giving rise to the changes in nuclear morphology and chromatin organization that were observed, and the role of the interactions between Ca^2+^ and cAMP in the process (Fig. 4a). Notably, the nuclear morphology of SRBW nanomechanostimulated cells that were prior treated with cytochalasin D—an inhibitor of actin polymerization—remained predominantly unaltered (Fig. 4b and Fig. S9 (Supporting Information)). A similar observation was noted when intra-cellular Ca^2+^ was chelated using BAPTA-AM (Fig. 4c, and Figs. S10a and S11 (Supporting Information)). Together, these results reveal the critical role of actin in modulating the organization and morphology of the nuclear periphery affecting the inner nuclear membrane Lamin organization: the interruption of the signal relay to the nucleus in the absence of the actin stress fibers, either due to disruption of actin polymerization or due to a lack of elevated intracellular Ca^2+^ levels, hints at the primary role of the actin cytoskeleton in transmitting extracellular signals from the plasma membrane to the nucleus via Nesprin proteins of the LINC complex, consistent with that previously reported [4, 17, 42, 60, 61, 62, 63, 64].

**Figure 4:**
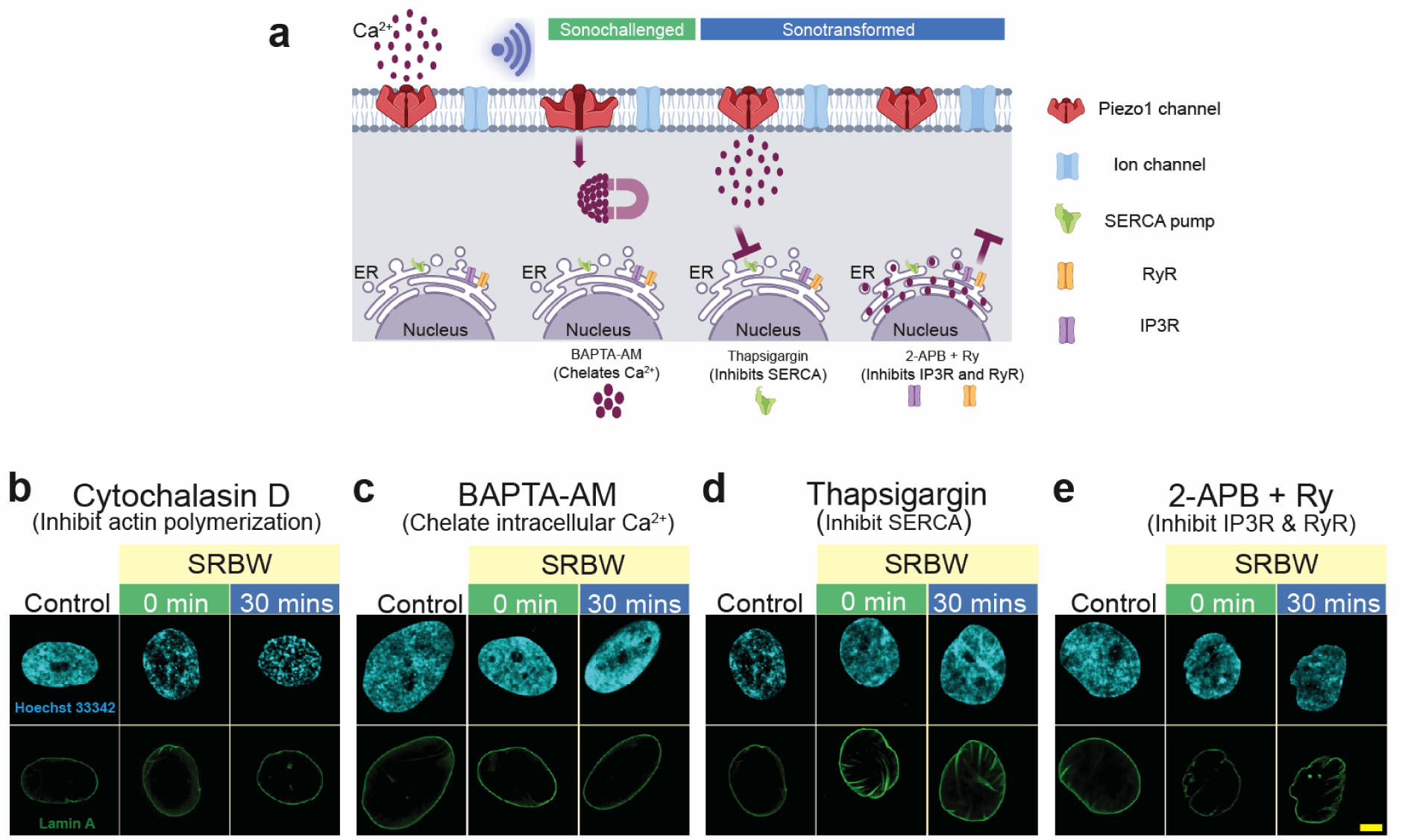
Actin polymerization regulates nuclear morphology upon SRBW nanomechanostimulation. (a) Schematic illustration showing the Ca^2+^ mobilization pathway in the presence of the various Ca^2+^ chelators and inhibitors: BAPTA-AM, a chelator of cytosolic Ca^2+^, thapsigargin, which blocks the SERCA pump to prevent Ca^2+^ influx into the ER, and, 2-APB and ryanodine (Ry), which impede the efflux of Ca^2+^ from the ER back into the cytosol. (b–e) Representative confocal images of Lamin A immunofluorescence (displayed in green) in the presence of (b) cytochalasin D, (c) BAPTA-AM, (d) thapsigargin, and, (e) 2-APB and ryanodine (Ry), at different post-exposure incubation compared to that for control (unstimulated) cells; all images were acquired at 100× magnification and the scale bars denote lengths of 5 µm.

In contrast, the nuclear morphological changes induced by the SRBW nanomechanostimulation were unaffected when elevation in intracellular cAMP levels were obstructed, either through inhibition of the sarco/endoplasmic reticulum calcium ATPase (SERCA) pump with thapsigargin to prevent influx of cytosolic Ca^2+^ into the ER (Fig. 4d, and Figs. S10a and S11 (Supporting Information)), or when the ER Ca^2+^ channels—IP3R and RyR—were blocked with 2-aminoethyl diphenylborinate (2-APB) and ryanodine (Fig. 4e, and Figs. S10b and S11 (Supporting Information)). We also note in these cases that the nuclear membrane maintained its wrinkled appearance up to 1 hr post-exposure, beyond the 30 min time point at which recovery began to occur in the uninhibited case. Both effects—be it the accumulation of cytosolic Ca^2+^ by blocking its influx into the ER through SERCA, or arresting the efflux of Ca^2+^ from the ER that is responsible for triggering cAMP signaling—led to the sustained presence of the actin stress fibers in the cytoskeletal network, thus delaying recovery of the nucleus from its deformation.

Taken together, these findings suggest that the SRBW mechanostimulation elicits a coordinated interplay in which the Piezo1-driven Ca^2+^ influx not only initiates a cAMP-dependent epigenetic program but also engages Rho/ROCK signaling in parallel by stabilizing actin-based force transmission. In particular, they highlight the centrality of the Ca^2+^ spatiotemporal dynamics (which, in turn, dictates the regulation of cAMP) in determining the nuclear morphological response to extracellular cues. The initial elevation of cytosolic Ca^2+^ during the sonochallenge phase activates the Rho/ROCK pathway in the cytoskeletal network to induce changes in nuclear morphology, mediated by the actin stress fibers, as a mechanism to provide parallel cytoskeletal reinforcement to the Ca^2+^–cAMP orchestrated chromatin remodeling. Mobilization of cytosolic Ca^2+^ into the ER, and the subsequent efflux of ER Ca^2+^ back into the cytosol through the CICR mechanism to trigger an increase in cytosolic cAMP levels in the sonotransformation phase, then initiates Epac/Rap1 signaling to counteract the Ca^2+^-triggered Rho/ROCK pathway and hence accommodate relaxation of the nuclear morphology back to its ground state. This bidirectional crosstalk between Ca^2+^ and cAMP therefore serves as a feedback mechanism from the nucleus as a means for maintaining cellular homeostasis [24, 37, 52, 53, 54, 55, 65, 66, 67, 68].

Unlike the nuclear morphological changes described above, the reorganization of the chromatin marks appear to remain unaffected when actin polymerization was disrupted by cytochalasin D (Fig. 5a,e). These results show that the actin cytoskeleton dynamics triggered by the SRBW nanomechanostimulation—which we observed previously to play a central role in the nuclear morphological response—do not interfere with the regulatory aspects of the chromatin structure observed. It therefore suggests that force transduction through the LINC complex that anchors the cytoskeletal network to the nuclear membrane is only one of the players relaying the extracellular cues to the nucleus. Instead, we observe the primary role of the spatiotemporal intracellular Ca^2+^ dynamics to be the dominant mechanism in driving chromatin fluctuations, as confirmed by the suppression of the histone chromatin marks observed (i.e., H3K27me3, H3K27ac and H3K9me3) upon removal of cytosolic Ca^2+^ with BAPTA-AM (Fig. 5b,f) or with inhibition of RyR and IP3R (Fig. 5d,h). Inhibiting Ca^2+^ influx into the ER with thapisgargin, on the other hand, can be seen to delay normalization of these histone markers (Fig. 5c,g).

**Figure 5:**
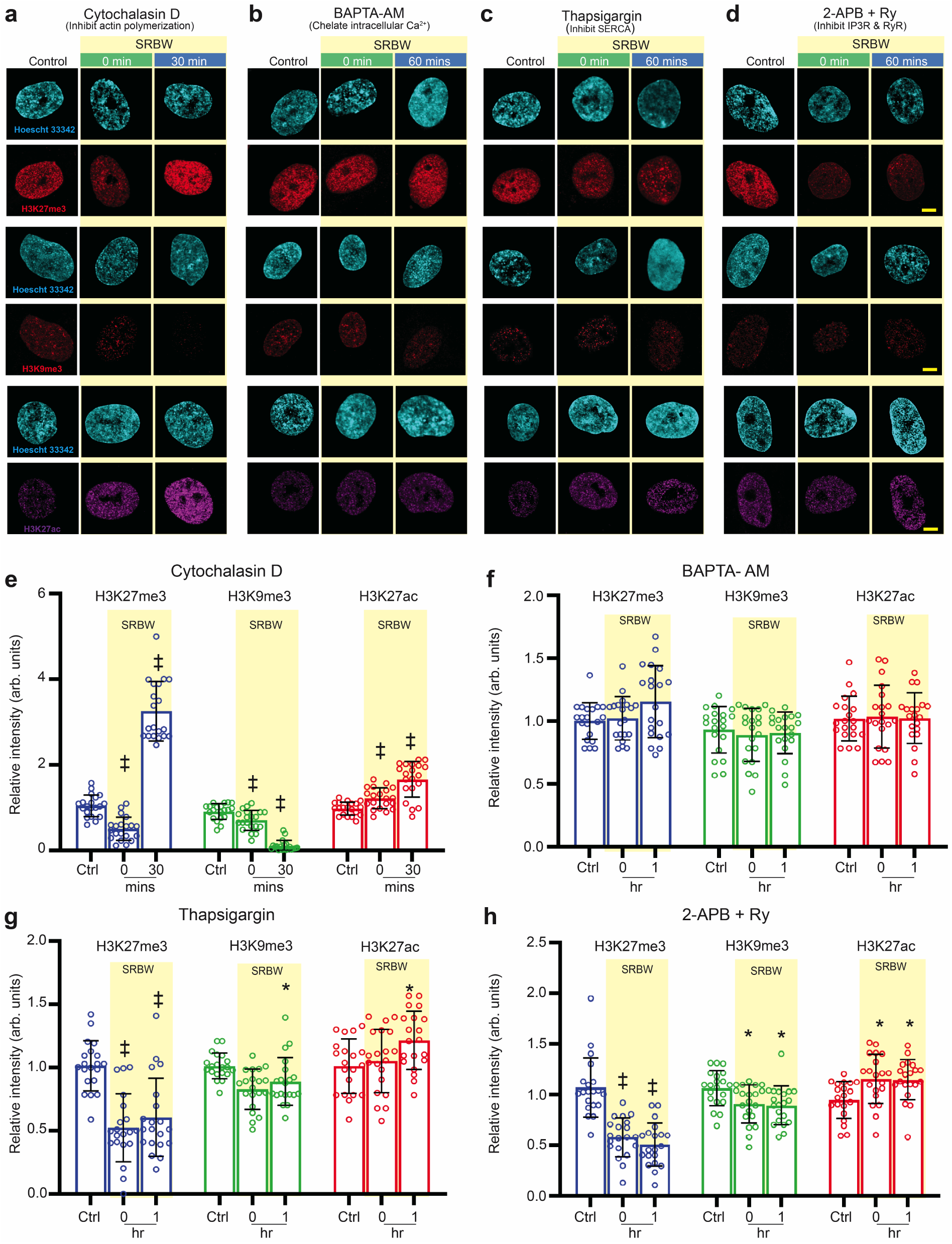
Ca^2+^ and cAMP dynamics regulate mechanoresponsive histone fluctuations in hMSCs. (a–d) Representative confocal microscopy images of the nucleus (stained with Hoechst 33342 and displayed in cyan) together with H3K27me3 (displayed in red), H3K9me3 (displayed in red) and H3K27ac (displayed in purple) immunofluorescence in the presence of (a) cytochalasin D, (b) BAPTA-AM, (c) thapsigargin, and, (d) 2-APB and ryanodine (Ry), at 0 and 30 mins or 1 hr post-exposure incubation following SRBW nanomechanostimulation, compared to that in the control (unstimulated) cells; all images were acquired at 100× magnification and the scale bars denote lengths of 5 µm. (e–h) Relative immunofluorescence of H3K27me3, H3K9me3 and H3K27ac in the SRBW mechanostimulated cells at 0 and 30 mins or 1 hr post-exposure incubation compared to that in the control (unstimulated) cells (Ctrl). Data are represented in terms of the mean value ± the standard error over multiple runs (*n* = 20 cells/condition, visualized from four independent experiments); ∗, † and ‡ indicate statistically significant differences with *p <* 0.05, *p <* 0.001 and *p <* 0.0001, respectively.

Disrupting actin polymerisation with cytochalasin D also had little effect on the SRBW-induced dynamics of RNAPII-S5p and the CBs, as can be seen in Fig. 6a. Chelating cytosolic Ca^2+^ with BAPTA-AM can be observed to prevent changes to the RNAPII-S5p and CB morphologies induced by the SRBW nanomechanostimulation (Fig. 6b). Additionally, maintaining elevated cytosolic Ca^2+^ levels while inhibiting intracellular cAMP by blocking the SERCA pump with thapsigargin led to the suppression of both RNAPII-S5p and CB cluster formation (Fig. 6c). We further noticed that inhibiting cAMP by blocking the RyR and IP3R channels prevented the formation of CB clusters and RNAPII-S5p condensation (Fig. 6d). Taken together, these results indicate that it is the Ca^2+^ spatiotemporal dynamics and the resulting cAMP signaling, rather than the actin cytoskeleton, that are the key mediators that predominantly guide the epigenetic modifications. Of particular interest is the paramount significance of the dynamics of second messenger Ca^2+^ and cAMP in regulating the chromatin fluctuations, and consequently, the transcription processes associated with nuclear mechanotransduction. This process, particularly its feedback response, constitutes a multifaceted molecular switch that influences the downstream fate of the cell, such as a return to homeostasis or a transition towards differentiation [69, 70].

**Figure 6:**
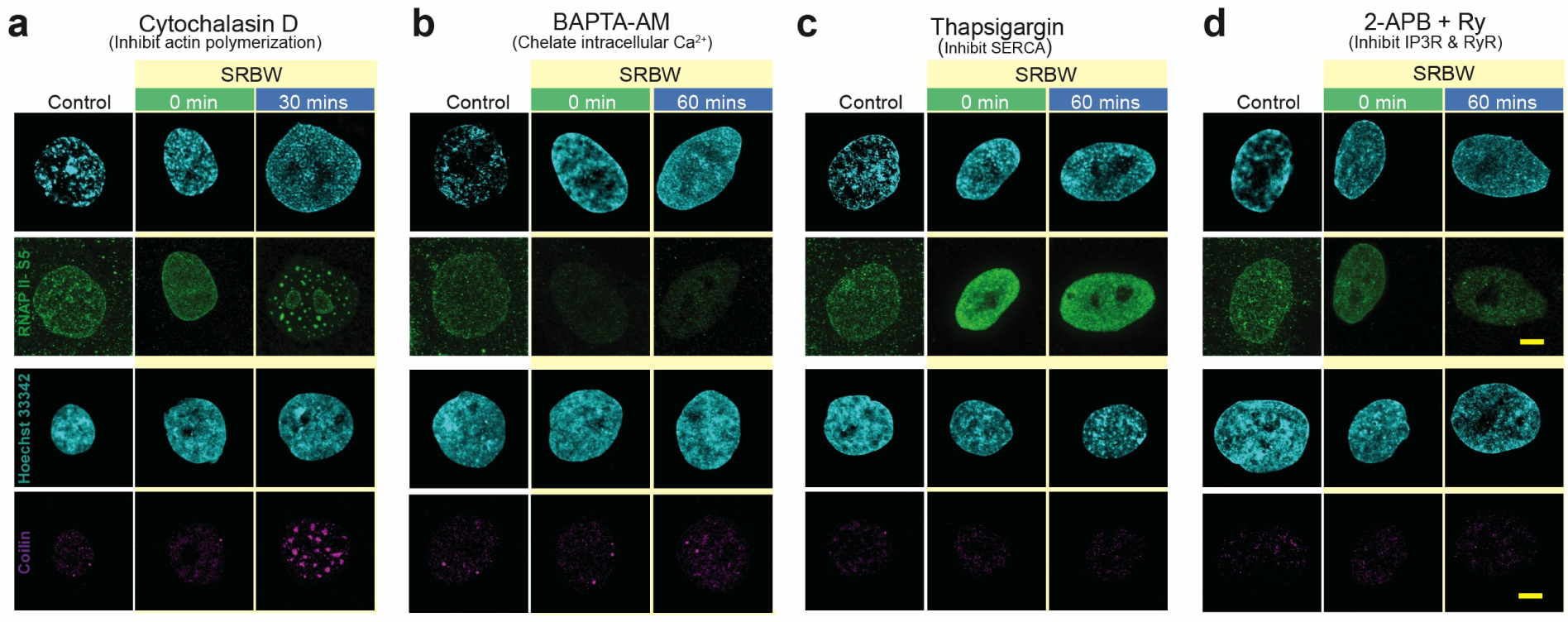
Nuclear body size fluctuations controlled by Ca^2+^ and cAMP dynamics. Representative confocal microscopy images of the nucleus (stained with Hoechst 33342 and displayed in cyan) together with RNAPII-S5p (displayed in green) and Coilin (displayed in purple) in the presence of (a) cytochalasin D, (b) BAPTA-AM, (c) thapsigargin, and, (d) 2-APB and ryanodine (Ry), at 0 and 30 mins or 1 hr post-exposure incubation following SRBW nanomechanostimulation, compared to that in the control (unstimulated) cells. All images were acquired at 100× magnification and the scale bars denote lengths of 5 µm.

### Downstream Cell Fate Regulation

We have previously reported that subjecting hMSCs to SRBW nanomechanostimulation 10 mins a day for 5 days led to the onset in osteogenic lineage commitment as early as 3 days [39]—without the need for osteogenic factors (see, also Fig. S12 (Supporting Information)). This is in distinct contrast, for example, with continuous mechanostimulation at kHz frequencies over 21 days, where commitment towards osteogenic differentiation was only observed after 5–7 days [71, 72]. The mechanism that enables such early osteogenic induction, however, has never been explicated to date. The data presented until now provide compelling correlative evidence that the Ca^2+^ and cAMP spatiotemporal dynamics induced by the SRBW nanomechanostimulation can provide a strong ‘mechanopriming’ effect that can initiate down-stream signaling cascades that drive the cells toward a specific fate. In order to demonstrate the mechanistic link between the Ca^2+^ and cAMP spatiotemporal dynamics, chromatin fluctuations and induced differentiation, we repeated the differentiation experiment in the presence of specific inhibitors. The results in Fig. 7a–e together with Fig. S13 (Supporting Information) show that inhibition of the Ca^2+^ and cAMP spatiotemporal dynamics downregulates specific osteogenic markers at the RNA as well as at protein level.

**Figure 7:**
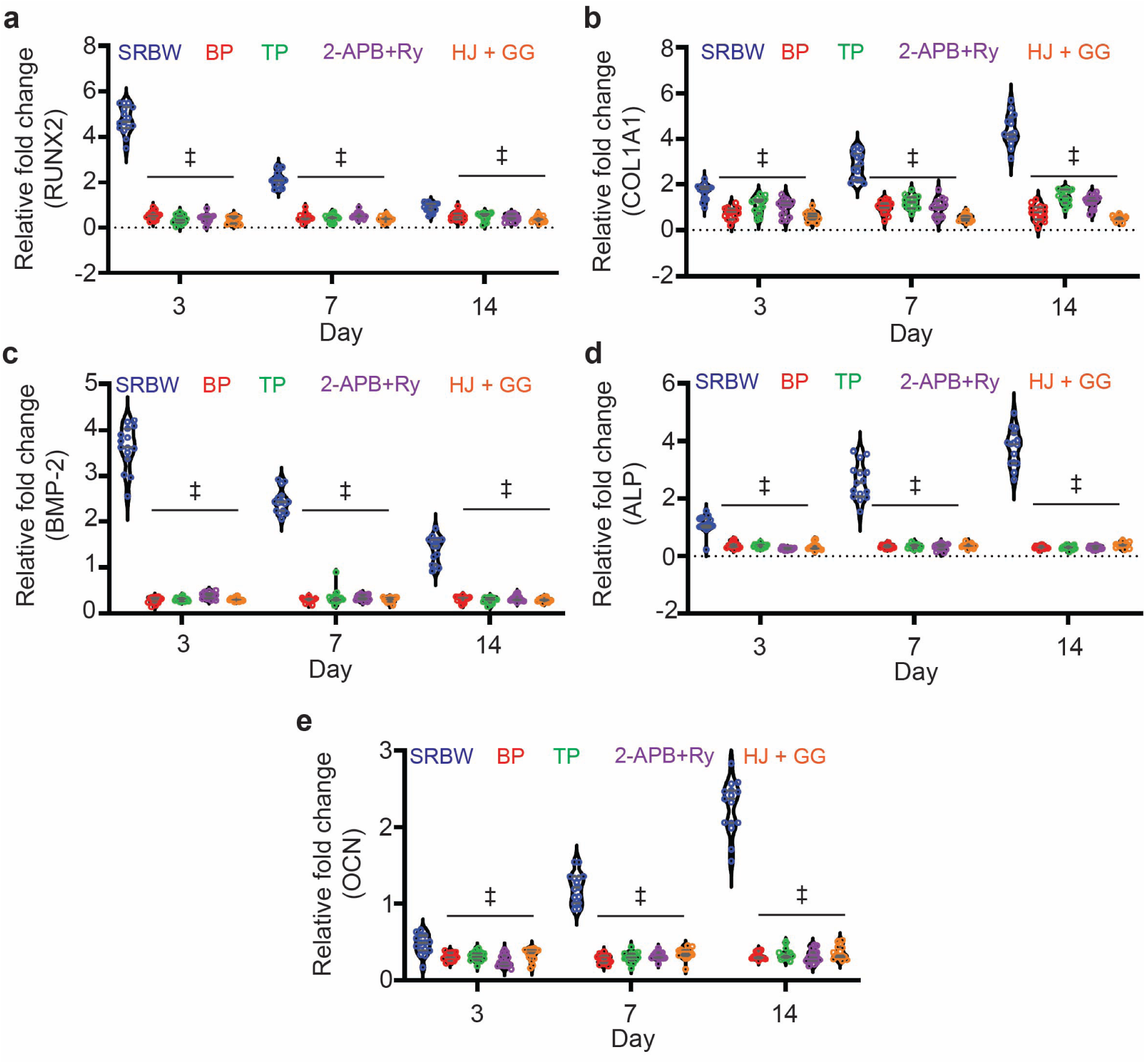
Early osteogenic lineage commitment in SRBW nanomechanostimulated hMSCs. Real-time quantitative polymerase chain reaction (RT-qPCR) analysis of (a) RUNX2 (runt homology domain transcription factor 2), (b) COL1A1 (collagen type I alpha 1), (c) BMP-2 (bone morphogenetic protein-2), (d) alkaline phosphatase (ALP), and, (e) osteocalcin (OCN), showing the mRNA relative fold change (against that of unstimulated cells in osteogenic media; i.e., the positive control) at different time points obtained for the SRBW nanomechanostimulated hMSCs in basal media, in the presence of the intracellular Ca^2+^ chelator BAPTA-AM (BP), 2-APB + Ry inhibiting increases in intracellular cAMP, an Epac1 inhibitor HJCO197 (HJ), and, a Rap1 inhibitor GGTI-298 (GG). The data represent the mean value ± the standard error over *n* = 15 experiments in triplicate runs; ‡ indicates statistically significant differences with *p <* 0.0001.

Interestingly, interfering the histone deacetylase activity through inhibition with sodium butyrate (NaB) can also be seen to suppress differentiation, therefore providing evidence for the crosstalk between the Ca^2+^–cAMP signaling and the observed epigenetic modifications (Figs. S14, S15a,b and S16 (Supporting Information)). Similarly, repeating the experiments with UNC1999—an EZH1/2 methyltransferase inhibitor that blocks PRC2 complex catalytic activity towards H3K27me3 deposition—can be seen to affect the osteogenic differentiation (Figs. S14, S15c, S16 and S17 (Supporting Information)). These results imply that a correct fluctuation of the chromatin marks in intensity and time is necessary to fulfill the correct epigenetic signatures associated with the osteogenic differentiation. Taken altogether, they confirm that the histone modifications arising due to the SRBW nanomechanostimulation are not merely correlative but necessary downstream effectors of Ca^2+^ and cAMP signaling associated with the mechanopriming process, therefore directly linking the SRBW-induced nuclear epigenetic rermodeling to functional osteogenic outcomes.

## Conclusions

Cells navigate a complex network of extracellular signals using precise signaling and regulatory mechanisms. To maintain cellular homeostasis and ensure proper function in both health and disease, it is essential to balance activation and inhibition, integrate multiple pathways, and adapt to external cues. The complex mechanotransduction pathways by which cells respond to persistent extracellular signals that alter their fate and function, however, has yet to be fully understood.

Through the use of high frequency nanomechanostimulation that allowed for spatiotemporal modulation of intracellular Ca^2+^ within the cell, we show that the cellular response to such extracellular mechanical cues, in its attempt to regain homeostasis, is dictated by the intricate spatiotemporal dynamics of these second messengers. Unlike the prevalent view that suggests the actin cytoskeleton as the direct and sole intermediary in relaying extracellular mechanical cues from the membrane to the nucleus via the LINC complex to exact epigenomic changes in the cell, we have found only changes in nuclear morphology to depend on such cytoskeleton transmission, and that this, in turn, is influenced by the biochemical signaling associated with the intracellular Ca^2+^ and cAMP dynamics. On the contrary, we have discovered the cells’ epigenomic response to be decoupled, at least directly, from the cytoskeletal/nuclear tether, and that it is instead mediated predominantly by Ca^2+^ dynamics and the secondary cAMP signaling it activates. In particular, the intricate bidirectional balance between these two second messengers— the latent cAMP-triggered Epac/Rap1 pathway following Ca^2+^-induced Rho/ROCK activation upon the mechanostimulation challenge—is central to driving the resultant modifications in the chromatin landscape and the activation of the relevant transcription machinery in the process of its recovery back towards a homeostatic state.

That the SRBW nanomechanostimulation induces dynamic, temporally-coordinated RNAPII-S5p clustering and CB formation, which occur in parallel with chromatin fluctuations, are consistent with a functional adaptive process in response to the mechanostimuli and seem to fit quite well with another study where a similar dynamic transition in RNAPII-S5p clustering follows a conserved condensation–dispersal when stem cells differentiate [70]. In line with these observations, it is possible that the RNAPII-S5p clustering may reflect the process of cellular transition towards a more differentiated state triggered by the SRBW nanomechanostimulation. As such, we hypothesize that the RNAPII-S5p and CB clustering reflects a nuclear sensing mechanism that integrates upstream Ca^2+^–cAMP signaling with transcriptional and epigenetic programs, rather than reflecting conformational instability of Coilin or RNAPII.

Along these lines, we show this conditioning in response to persistent stimuli gives rise to a mechanopriming effect that explains why they were observed to commit to an osteogenic lineage much earlier than has been typically observed. This discovery thus reframes the nuclear response to mechanical cues—not just as passive adaptation of the cell in response to mechanical stress, but as a dynamic regulatory system that actively drives chromatin fluctuations and transcriptional priming to effectively direct downstream cell fate.

## Materials and Methods

### Materials

Bone marrow derived human mesenchymal stem cells (hMSCs) were obtained from the American Type Culture Collection (PCS-500-012™; ATCC, Manassas, VA, USA). To minimize donor-specific variability and hence ensure reproducibility, cells from three independent healthy adult donors were used. All experiments were performed with cells between passages 3 and 5.

Mesenchymal stem cell basal medium (MSCBM™), mesenchymal stem cell growth BulletKit™ medium (MSCGM™) and osteogenic differentiation BulletKit™ medium were procured from Lonza Pty. Ltd. (Mount Waverley, VIC, Australia). Sodium chloride, potassium chloride, magnesium chloride, disodium hydrogen phosphate, sodium bicarbonate, methanol, ethanol, isopropanol, hydrogen chloride (HCl), ammonium hydroxide, acetic acid, glucose, silicon oil, glycerol, ethylenediaminetetraacetic acid (EDTA), 3-isobutyl-1-methylxanthine (IBMX), formaldehyde, dimethylsulfoxide (DMSO), *β*-mercaptoethanol, alizarin red S, trypan blue, bromophenol blue, sodium dodecyl sulfate (SDS), Tween 20, TRIZMA™ base, nitrocellulose membrane (0.45 µm), protease inhibitor cocktail tablets, skimmed milk powder, polyacrylamide gel, ruthenium red (RR) and 1-isopropyl-6-(6-(4-isopropylpiperazin-1-yl)pyridin-3-yl)-N-((6-methyl-2-oxo-4-propyl-1,2-dihydropyridin-3-yl)methyl)-1H-indazole-4-carboxamide (UNC1999) were acquired from Sigma-Aldrich Pty. Ltd. (Bayswater, VIC, Australia). 2-aminoethyl diphenylborinate (2-APB), ryanodine (Ry), 4-cyclopentyl-2-[[(2,5-dimethylphenyl)methyl]thio]-1,6-dihydro-6-oxo-5-pyrimidinecarbonitrile (HJC0197), N-[[4-(2-(R)-amino-3-mercaptopropyl)amino]-2-naphthylbenzoyl]leucine methyl ester trifluoroacetate salt hydrate (GGTI-298) were obtained from R&D systems (Minneapolis, MN, USA); and, 3-(4,5-dimethylthiazol-2-yl)-2,5-diphenyltetrazolium bromide (MTT), TRiZOL™ reagent, bovine serum albumin (BSA), fetal bovine serum (FBS), Dulbecco’s Modified Eagle Medium (DMEM), acrylamide/bisacrylamide Solution (40%, 37.5:1), 3,3’,5,5’’-tetramethylbenzidine (TMB), Fura-2 acetoxymethyl ester (Fura-2AM),1,2-bis(2-aminophenoxy) ethane-N,N,N’,N’-tetraacetic acid tetrakis(acetoxymethyl ester) (BAPTA-AM), thapsigargin, cytochalasin D, sodium butyrate (NaB), ActinRed™ 555 ReadyProbes™, NucBlue™ Live ReadyProbes™, anti-RUNX2 antibody, Pierce ECL Western blotting detection reagent, bicinchoninic acid (BCA) protein assay kit, T25 cell culture flasks, Nunc™ 8-well plate, and, Nunc™ Lab-Tek™ II Chamber Slide and Chambered Coverglass from Thermo Fischer Scientific Pty. Ltd. (Scoresby, VIC, Australia); whereas the LunaScript® RT SuperMix kit and Luna® Universal qPCR Master Mix were obtained from New England BioLabs (Notting Hill, VIC, Australia). FluoroDish cell culture plates were purchased from World Precision Instruments (WPI) Ltd. (Hertfordshire, England). Cyclic AMP XP™ assay, radio-immunoprecipitation assay (RIPA) buffer, anti-GAPDH antibody, anti-COL1A1 antibody, acetylated histone H3 ELISA kit, anti-Lamin A antibody, anti-H3K9me3 antibody, anti-H3K27me3 antibody, anti-H3K27ac antibody, anti-phospho-H2A.X antibody, anti-Coilin antibody, anti-RNAPII Ser5 antibody, anti-rabbit and anti-mouse IgG (H+L) F(ab’)2 Fragment (Alexa Fluor 488 conjugate), anti-rabbit IgG (H+L) F(ab’)2 Fragment (Alexa Fluor® 555 Conjugate) and F(ab’)2 Fragment (Alexa Fluor® 647 Conjugate), on the other hand, were obtained from Cell Signaling Technology Inc. (Danvers, MA, USA).

The following primers used for RT-qPCR analysis were acquired from Integrated DNA Technologies Inc. (Coralville, IA, USA):

*β*-actin (forward): 5’-TGACGGGGTCACCCACACTGTGCCCAT-3’,
*β*-actin (reverse): 5’-CTAGAAGCATTTGCGGTGGACGATGGA GGG-3’,
ALP (forward): 5’-ATGAAGGAAAAGCCAAGCAG-3’,
ALP (reverse): 5’-CCACCAAATGTGAAGACGTG-3’,
COL1A1 (forward): 5’-ACATGTTCAGCTTTGTGGACC-3’,
COL1A1 (reverse): 5’-TGATTGGTGGGATGTCTTCGT-3’,
BMP-2 (forward): 5’-ATGGATTCGTGGTGGAAGTG-3’,
BMP-2 (reverse): 5’-GTGGAGTTCAGATGATCAGCG-3’,
OCN (forward): 5’-GACTGTGACGAGTTGGCTGA-3’,
OCN (reverse): 5’-CTGGAGAGGAGCAGAACTGG-3’,
RUNX2 (forward): 5’-AAGTGCGGTGCAAACTTTCTT-3’,
RUNX2 (reverse): 5’-TCTCGGTGGCTGGTAGTGA-3’,
PP1A (forward): 5’-CGGGTCCTGGCATCTTGT-3’,
PP1A (reverse): 5’-CAGTCTTGGCAGTGCAGATGA-3’.

### Device fabrication

The SRBW devices illustrated in Fig. 1a and Fig. S1 (Supporting Information) consisted of 40 alternating fingers, each 11 mm wide and 66 nm thick, arranged as a straight interdigitated aluminum transducer (IDT) electrode in a basic full-width interleaved design on 500-µm-thick 127.86^◦^ *Y* –*X* lithium niobate (LiNbO_3_) single crystal piezoelectric substrates (Roditi Ltd., London, UK). The IDT was patterned on a 33-nm-thick chromium adhesion layer using sputter deposition and standard ultraviolet (UV) photolithography. The width and separation of the IDT fingers (*λ*/4) corresponds to the SRBW wavelength *λ*, which is set to be 398 µm, which, in turn, determines the resonant frequency of the device *f* = *c/λ* = 10 MHz, wherein *c* is the phase speed in LiNbO_3_. To produce the SRBW, an alternating electrical signal at the resonant frequency was applied to the IDT using a signal generator (SML01, Rhode & Schwarz Pty. Ltd., North Ryde, NSW, Australia) and amplifier (10W1000C, Amplifier Research, Souderton, PA, USA). The acoustic wave energy was then transmitted to a glass-bottom square-well chamber slide (0.15 mm thickness) containing the adherent cells through a thin layer of silicon oil with viscosity and density of 45–55 cP and 0.963 g/mL at 25^◦^C, respectively.

### Cell culture and mechanostimulation

The hMSCs were grown in DMEM with 1% penicillin–streptomycin and 10% FBS in a humidified incubator at 37^◦^C and 5% CO_2_ until they covered 80–90% of a standard T25 flask. They were then detached using 0.05% trypsin–EDTA and transferred to 8-well plates or FluoroDish cell culture plates at a seeding density of 3,000 cells per well before being incubated for 48 hrs in DMEM to ensure proper adhesion. The cells in the well-plate were subsequently exposed to the 10 MHz SRBW irradiation at an input power of 2.5 W for 10 mins. Control samples consisted of cells seeded in DMEM with 1% penicillin–streptomycin and 10% FBS at the same density and incubated for the same time period, but without any vibrational excitation. The cells were then fixed at 0, 5, 15 and 30 mins, and 1, 2, 4 and 8 hrs post-exposure incubation.

To facilitate osteogenic differentiation, the cells were cultured in a standard T25 flask until they achieved 80–90% confluency. At this point, they were detached using 0.05% trypsin– EDTA and replated into 8-well plates at a density of 3,000 cells per well, allowing for a 48 hr incubation in basal media (BM; *α*–MEM) to ensure adequate adhesion. After this incubation, the media was refreshed with new basal media. The cells in the well plates were then subjected to irradiation with the 10 MHz SRBW at an input power of 2.5 W for a duration of 10 mins, which represented a single treatment regimen (1×). Once the vibrational excitation ceased, the cells were left to incubate overnight at 37^◦^C. The 1× treatment regimen was applied continuously for the following four days. Following this five-day regimen, the cells underwent an additional incubation period of 17 days. Control samples for real-time quantitative polymerase chain reaction (RT-qPCR) consisted of cells plated in osteogenic media (OM) at the same density, which were incubated over the same timeframe but without any form of vibrational excitation.

The following inhibitor concentrations were used: actin polymerisation inhibitor (cytochalasin D): 5 µM; intracellular Ca^2+^ chelator (BAPTA-AM): 20 µM; SERCA ion channel blocker (thapsigargin): 300 nM; IP3R inhibitor (2-APB): 50 µM; RyR inhibitor (ryanodine): 15 µM; histone deacetylase inhibitor (NaB): 1.15 mM; EZH1/2 inhibitor (UNC1999): 200 nM; Piezo channel blocker (RR): 10 µM.

Experimental replicates were achieved by performing independent biological experiments using separate hMSCs cultures (biological replicates), typically seeded in individual wells atop separate SRBW devices. For each replicate, either new or previously used devices were employed. In the latter case, devices were sterilized following each use by thoroughly removing the silicon oil couplant and cleaning with ethanol and isopropanol, although we note that there is minimal risk of contamination since the cells were cultured on glass-bottom plates and were hence not in direct contact with the device. Each stimulation run was performed under identical parameters (frequency, amplitude, and duration), with multiple wells stimulated either in parallel or sequentially within the same session to ensure reproducibility (technical replicates). Unless otherwise stated, all experiments were conducted in triplicate, and data were pooled from at least three independent experiments to ensure statistical validity.

For the time-course experiments, two independent replicates were performed within the same experimental session using cells derived from three separate culture flasks. An additional two replicates were carried out in a separate session within 20 days of the initial experiments, under identical conditions. Across all sessions, stimulation parameters (frequency, amplitude, duration), cell seeding density, reagent concentrations and volumes were carefully maintained to ensure consistency and reproducibility.

### Immunofluorescence staining

The cells requiring fixation were washed three times in PBS and then incubated in 4% formaldehyde for 20 mins at room temperature. Following this, they were washed three more times in PBS. To block non-specific binding, the cells were then incubated in 1X 5% normal serum in Triton™ X-100 for 60 minutes. They were subsequently incubated overnight at 4^◦^C with primary antibodies: anti-Lamin A (1:2000), anti-H3K27me3 (1:800), anti-H3K27ac (1:200), anti-H3K9me3 (1:400), and either anti-RNAPII Ser5 or anti-Coilin (1:800) diluted using 1X PBS/0.3% Triton™ X-100. After three washes in PBS, the cells were incubated for 1 hr in the dark at room temperature with the secondary antibodies (anti-rabbit and anti-mouse IgG (H+L) F(ab’)2 Fragment (Alexa Fluor 488 conjugate); 1:1000), followed by three additional washes in PBS. Nuclei were counterstained using NucBlue™ Live ReadyProbes™ and actin filaments were stained with ActinRed™ 555 ReadyProbes™ before imaging, which was performed on an inverted confocal laser scanning microscope (FV3000; Olympus, Tokyo, Japan).

### Intracellular Ca^2+^

The fluorescent calcium indicator Fura-2AM was utilized to measure free cytosolic Ca^2+^ levels. The cells were incubated in 5 µmol/l Fura-2AM mixed with Opti-MEM™ reduced serum medium containing 2% (vol/vol) heat-inactivated FBS at 37^◦^C in the absence of light. Following a 1 hr incubation period, the medium containing extracellular Fura-2AM was replaced with fresh medium, and the cells were incubated for an additional 20 mins before subjecting the samples to irradiation with the SRBW. Fluorescence emission intensity was then measured at 510 nm using a spectrophotometric plate reader (CLARIOstar, BMG LabTech, Mornington, VIC, Australia) in individual wells, employing excitation wavelengths of 340 nm and 380 nm. The ratio of Fura-2AM fluorescence emission in response to the 340 nm and 380 nm excitation (340/380) was calculated and expressed as the fold change relative to the respective control (unstimulated) cells.

### cAMP

cAMP levels were assessed using a cyclic AMP XP™ assay. Initially, the cells underwent washing with Tyrode’s buffer (137 mM sodium chloride, 12 mM sodium bicarbonate, 5.5 mM glucose, 2 mM potassium chloride, 1 mM magnesium chloride, and 0.3 mM disodium hydrogen phosphate) at pH 7.4, which was supplemented with 100 µmol/l IBMX—a phosphodiesterase inhibitor—to prevent the breakdown of cAMP. The cells were subsequently treated with the SRBW and incubated at various time intervals, following which they were lysed in 100 µl of the lysis buffer provided by the kit. A total of 25 µl of the cell lysate and HRP-linked cAMP solution was placed into the designated cAMP assay microtiter plates and incubated at room temperature for 3 hrs. After incubation, the plate underwent four washing cycles, and the excess liquid was discarded. Then, 100 µl of 3,3’,5,5’’-tetramethylbenzidine (TMB) was added, and the solution was kept in the dark for 15 mins. The reaction was terminated by adding 100 µl of the provided STOP solution, and its absorbance was assessed at 450 nm with a spectrophotometric plate reader (CLARIOstar, BMG LabTech, Mornington, VIC, Australia). The average of three measurements for each treatment group was calculated from cells derived from each individual culture, allowing for the determination of cAMP concentrations based on a standard curve.

### RNA isolation and real-time quantitative polymerase chain reaction (RT-qPCR)

RNA was extracted from control and SRBW-treated cells using TRiZOL™. Initially, the cells were homogenized in TRiZOL™, and chloroform was added to the mixture. After centrifugation, the RNA-containing aqueous layer was separated. The RNA was then precipitated with isopropanol, washed with ethanol, and resuspended in RNase-free water containing 0.1 µM EDTA. The RNA concentration was measured using a UV spectrophotometer (NanoDrop™ One; Thermo Fisher Scientific, Waltham, MA, USA). Subsequently, cDNA synthesis was carried out using the LunaScript® RT SuperMix kit, and RT-qPCR was conducted with the Luna® Universal qPCR Master Mix along with the aforementioned designated primers.

### Acetylation quantification

Changes in histone H3 acetylation were assessed using an acetylated histone H3 ELISA kit following the manufacturer’s protocol. The hMSCs were divided into four groups: (1) unstimulated cells, (2) SRBW-stimulated cells, (3) unstimulated cells treated with NaB, and, (4) NaB-treated cells exposed to the SRBW. Cells were harvested at 0 min, 30 mins, 1 hr, 4 hrs, and 8 hrs post-exposure (with matching time points for unstimulated controls). Cell lysates were prepared, and equal amounts of protein were applied to ELISA plates pre-coated with capture antibody. After incubation with the detection antibody and the HRP-conjugated secondary antibody, the signal was developed with TMB substrate and measured at 450 nm using a plate reader (CLARIOstar, BMG LabTech, Mornington, VIC, Australia). Absorbance values were normalized to the unstimulated control to compare temporal changes in acetylation across experimental groups.

### Western blotting

After the stipulated incubation period post-exposure, cells were lysed in RIPA buffer supplemented with a protease inhibitor cocktail. Lysates were mixed with reducing SDS sample buffer (62.5 mM Tris–HCl, pH 6.8; 2% SDS; 25% glycerol; 0.01% bromophenol blue; and 5% freshly added *β*-mercaptoethanol) and heated at 95^◦^C for 5 min. Proteins were resolved on 12% polyacrylamide gels and electrotransferred to nitrocellulose membranes (60 mV; 45 mins (H3K27ac, H3K27me3 and GAPDH), 90 mins (RNAP)). Membranes were blocked for 1 hr in 5% skimmed milk prepared in TBST (20 mM Tris, 150 mM NaCl, 0.05% Tween-20) and then incubated overnight at 4^◦^C with primary antibodies (1:2000) and anti-rabbit secondary antibodies (1:50,000). Following washes, HRP-conjugated secondary antibodies were applied in 0.05% TBST at room temperature for 1 hr, and signals were developed using Pierce ECL reagents (2 mins, room temperature). Chemiluminescence was detected on an imager (LI-COR Biotechnology, Lincoln, NE, USA), with GAPDH serving as the loading control.

### Mineralization assessment

Mineralization was evaluated by alizarin red staining at day 16 post-seeding (17 days after the first SRBW mechanostimulation). Briefly, 0.2% alizarin red solution was prepared in distilled water and adjusted to pH 6.4 using ammonium hydroxide. Following the culture period, cells were washed thrice with PBS, rinsed in distilled water, and fixed in 100% ethanol for 15 mins. Samples were then incubated with the alizarin red solution for 1 hr at room temperature and imaged using an optical microscope (Eclipse TS 100; Nikon Instruments Inc., Melville, NY, USA). For quantification, stained wells were washed twice with PBS and incubated with leaching solution (20% methanol and 10% acetic acid in distilled water) under gentle agitation for 15 mins. Absorbance of the eluate was recorded at 450 nm using a plate reader (CLARIOstar, BMG LabTech, Mornington, VIC, Australia), with leaching solution as the blank. Values were normalized against the negative control (hMSCs grown in basal media (BM)).

### Image analysis

Nuclear morphometry (volume, area, thickness), fluorescent intensity and transcription hub characteristics (number and size in diameter) were measured using ImageJ (v1.54p; National Institutes of Health, Bethesda, MD, USA), whereas the nuclear shape descriptors were measured using the measurement module in the CellProfiler tool (v4.2.8; Broad Institute, Cambridge, MA, USA).

### Statistics

The data provided in this study were represented as the average ± the standard error. Where applicable, the data were assessed using ordinary one-way or two-way analysis of variance (ANOVA) or with a two-tailed, unpaired Student’s t-test.

## Acknowledgments

LYY acknowledges support from the Australian Research Council through Discovery Project grant DP210101720. CF was supported by the clinical research unit 5002 of the Deutsche Forschungsgemeinschaft (KFO5002).

## Author contributions

**Conceptualisation**: LAA and LYY; **Methodology**: LAA and CF; **Data curation**: LAA; **Formal analysis**: LAA, CF and LYY; **Investigation**: LAA, CF and LYY; **Visualization**: LAA, CF and BdR; **Supervision**: CF and LYY; **Writing – original draft**: LAA, CF and LYY; **Writing – review & editing**: all authors

## Competing interests

LAA and LYY are inventors on a patent application for the use of the method for differentiating stem cells (PCT/AU2023/050012). The remaining authors have no competing interests to declare.

## Data and materials availability

All data needed to evaluate the conclusions in the paper are present in the paper and/or the Supplementary Materials.

## Supplementary Materials

Figs. S1–S17

## Notes

### Competing Interest Statement

The authors have declared no competing interest.

### Summary of Updates

Title change; New supplementary figures added; Addition of new results for osteogenic differentiation, including additional protein-level characterisation and inhibitor studies

