## Supplementary Figures S1-S17 for "Sonoepigenetic Modification Mechanoprimes Early Osteogenic Commitment in Mesenchymal Stem Cells"

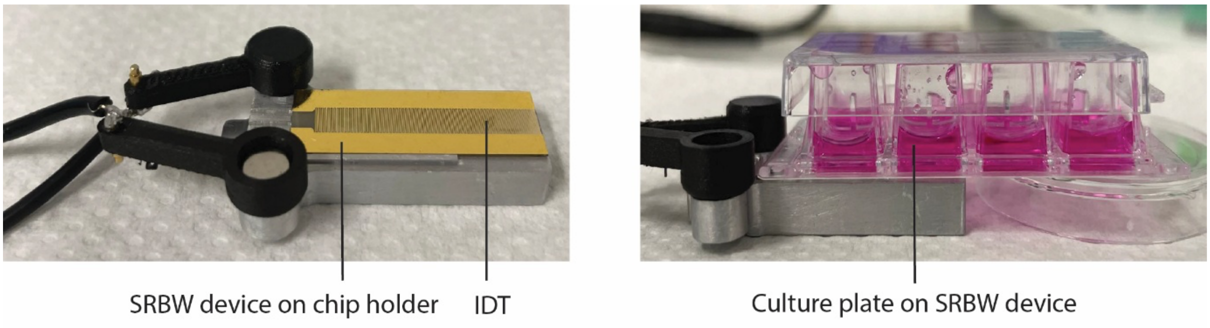


**Figure S1** Images depicting the experimental setup (not to scale). The SRBW is generated by applying a sinusoidal electrical signal to an interdigitated transducer (IDT) that has been photolithographically patterned onto a single crystal piezoelectric substrate ($LiNbO_3$)—shown in the left image—at its resonant frequency (10 MHz). As shown in the right image, the SRBW is then coupled through a thin layer of silicon oil into a glass-bottomed culture plate containing the adherent human mesenchymal stem cells (hMSCs).

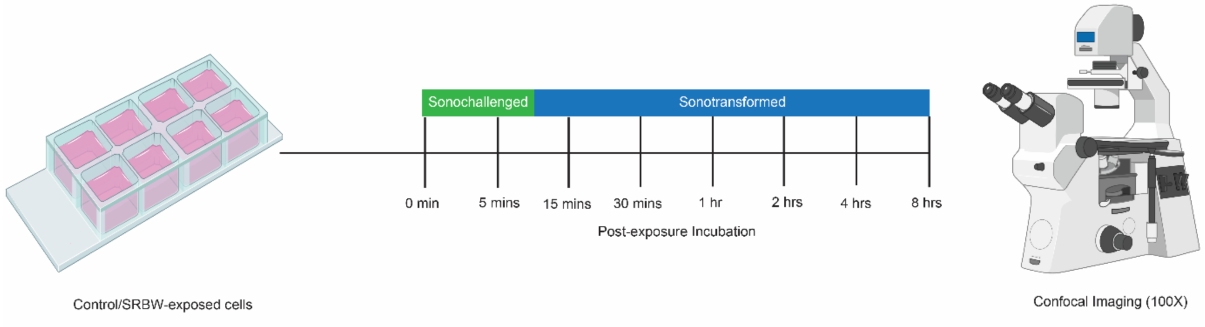


**Figure S2** Time points used for the nuclear response studies: the SRBW-exposed cells were fixed at the end of the 10 min nanomechanostimulation period at 0 hrs, and assessed subsequently at 5, 15 and 30 mins, and, 1, 2, 4 and 8 hrs post-exposure; these time points were selected to correlate with the relevant timescales associated with the intracellular $Ca^{2+}$ and cAMP dynamics, which also concurred with the characteristic times associated with the actin reorganization.

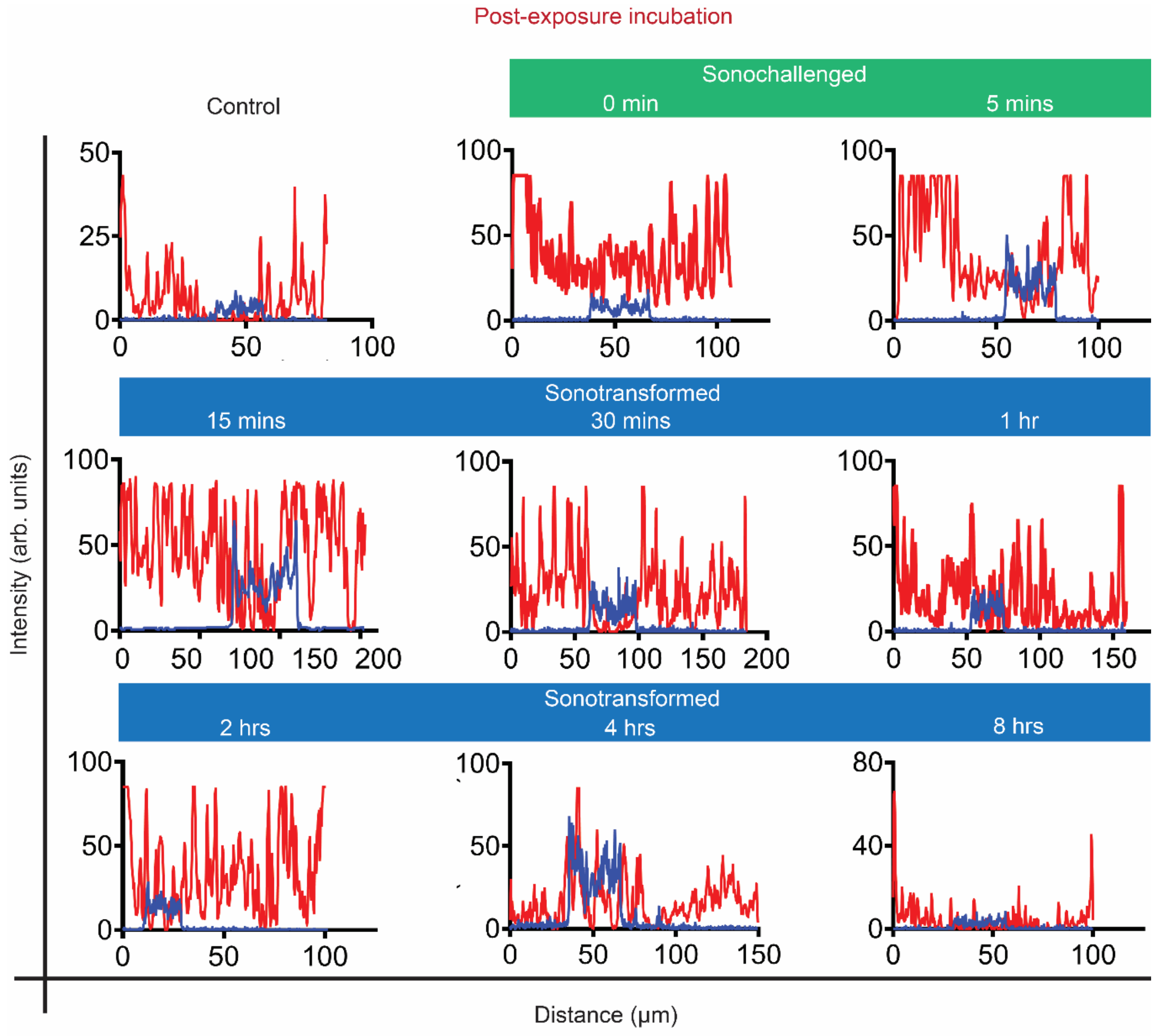


**Figure S3** Intensity profile along a line drawn across the nucleus for the actin (red) and nuclear (blue) fluorescence signals for representative cells exposed to the SRBW nanomechanostimulation at different post-exposure incubation times.

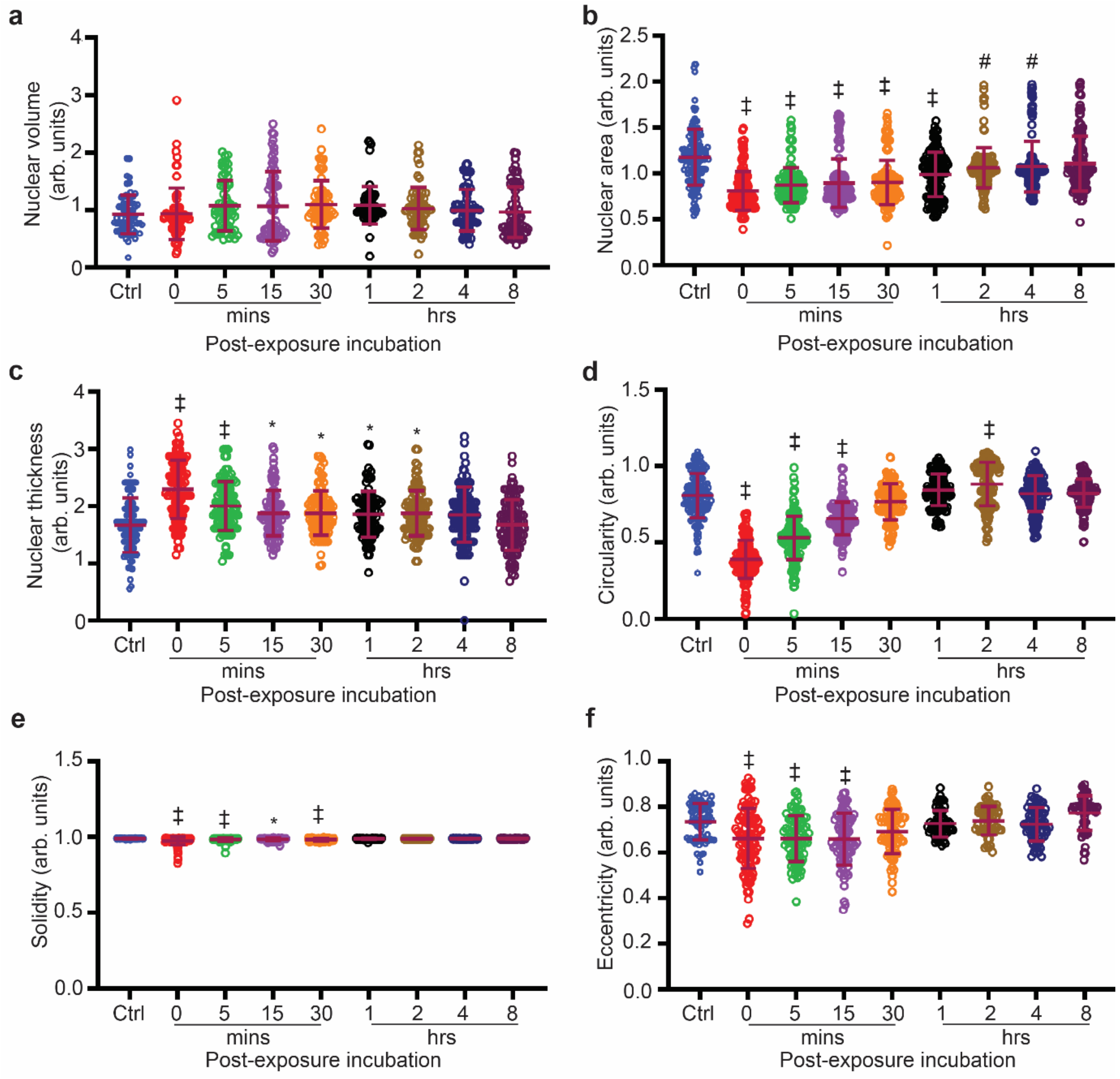


**Figure S4** Quantification of changes in the nuclear morphometry due to the SRBW nanomechanostimulation as a function of the post-exposure incubation time, relative to that of the control (unstimulated) cells. Nuclear (a) volume (measured in the *xyz*-space), (b) area (measured as a projection in the *xy*-plane), and, (c) thickness (i.e., the maximum length from the basal surface to the apical surface, measured as a projection of the three-dimensionally reconstructed nuclei onto the *xz*-plane), in addition to other nuclear shape factors, such as its (d) circularity, (e) solidity, and, (f) eccentricity. The data are represented in terms of the mean value ± the standard error over multiple runs ($n > 50$ cells/condition, visualized from four independent experiments); *, # and ‡ indicate statistically significant differences with $p < 0.05$, $p < 0.01$ and $p < 0.0001$, respectively.

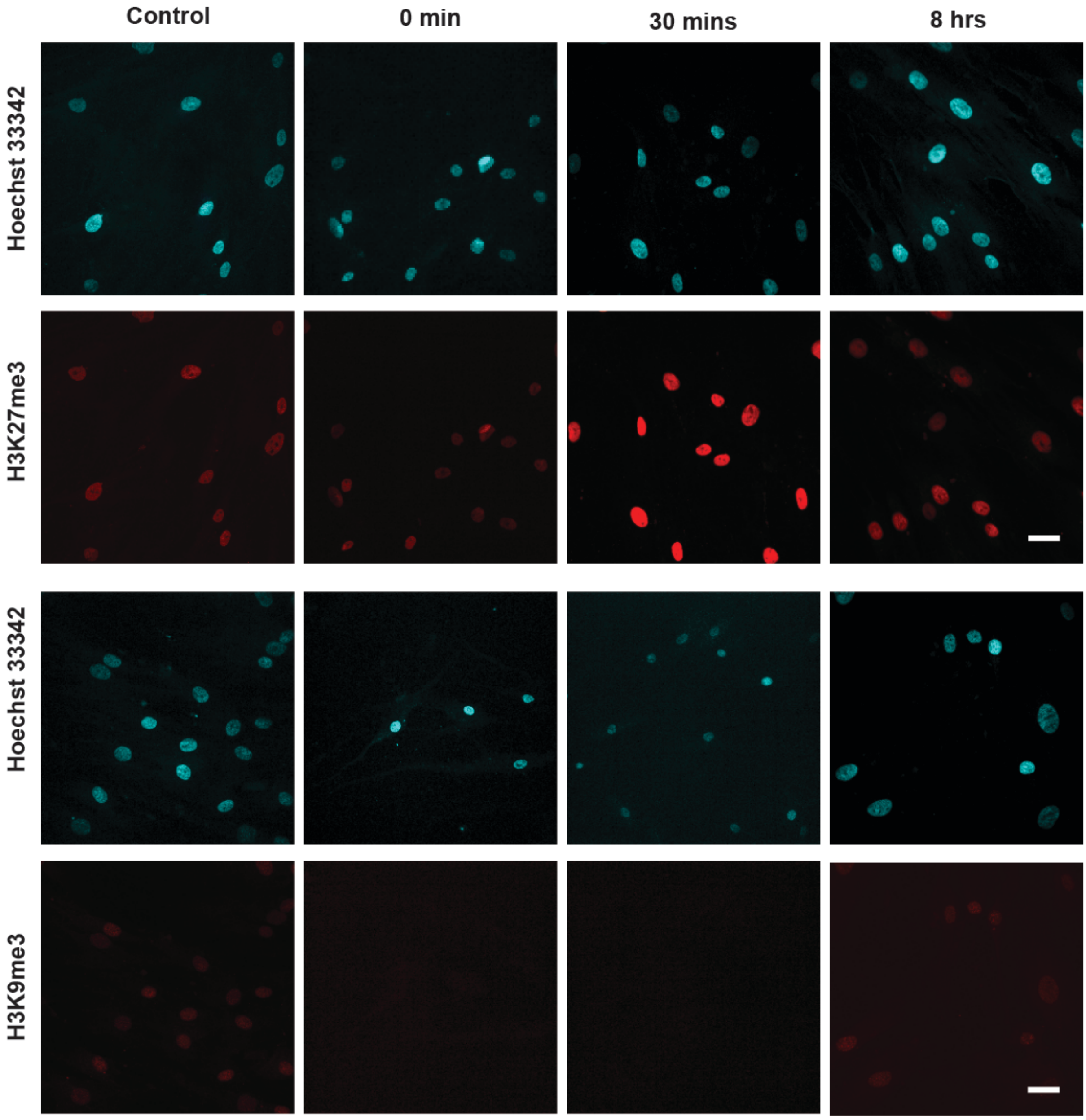


**Figure S5** Representative confocal microscopy images of the nucleus (stained with Hoechst 33342 and displayed in cyan) together with H3K27me3 and H3K9me3 (displayed in red) immunofluorescence following SRBW nanomechanostimulation of the hMSCs at different post-exposure incubation times, compared to those of the control (unstimulated) cells. The scale bars denote lengths of 40 µm.

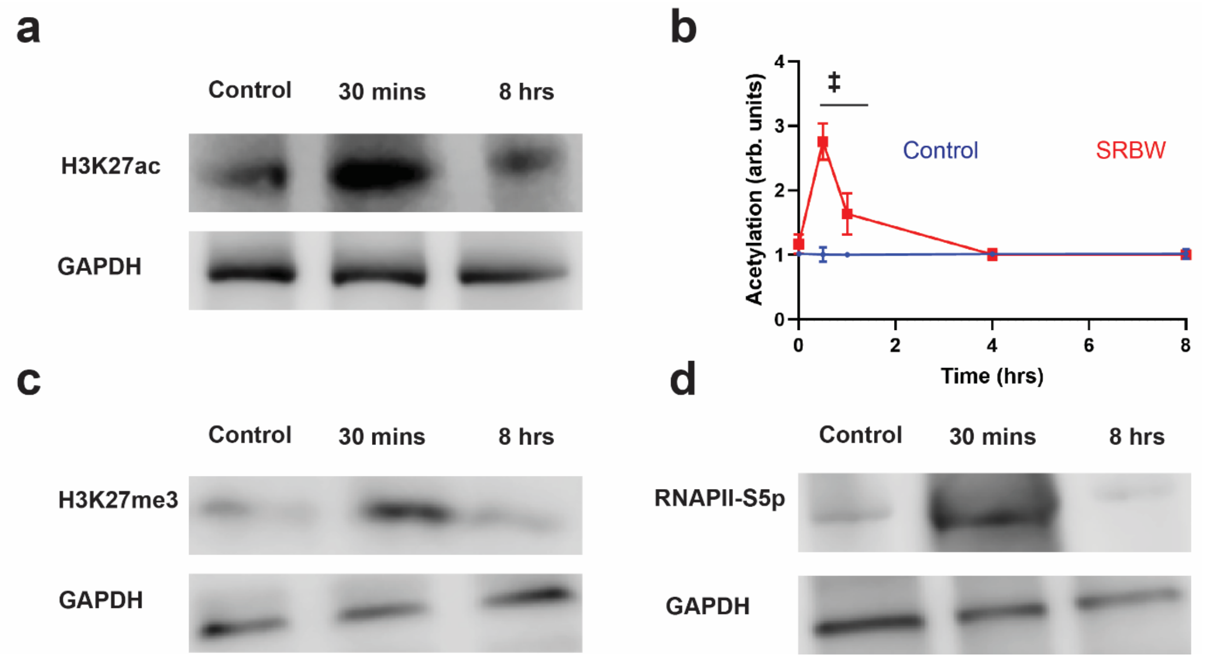


**Figure S6** (a,c,d) Western blots of (a) H3K27ac, (c) H3K27me3, and, (d) RNAPII-5Sp for unstimulated hMSCs in basal media (negative control) and SRBW nanomechanostimulated hMSCs for different post-exposure incubation periods, along with that of the loading control (GAPDH). (b) Quantification of H3 acetylation in hMSCs by ELISA with time for the case when the cells the unstimulated and SRBW nanomechanostimulated cells (normalized against the case of unstimulated cells); data are represented in terms of the mean value ± the standard error; ‡ indicates statistically significant differences with $p < 0.0001$.

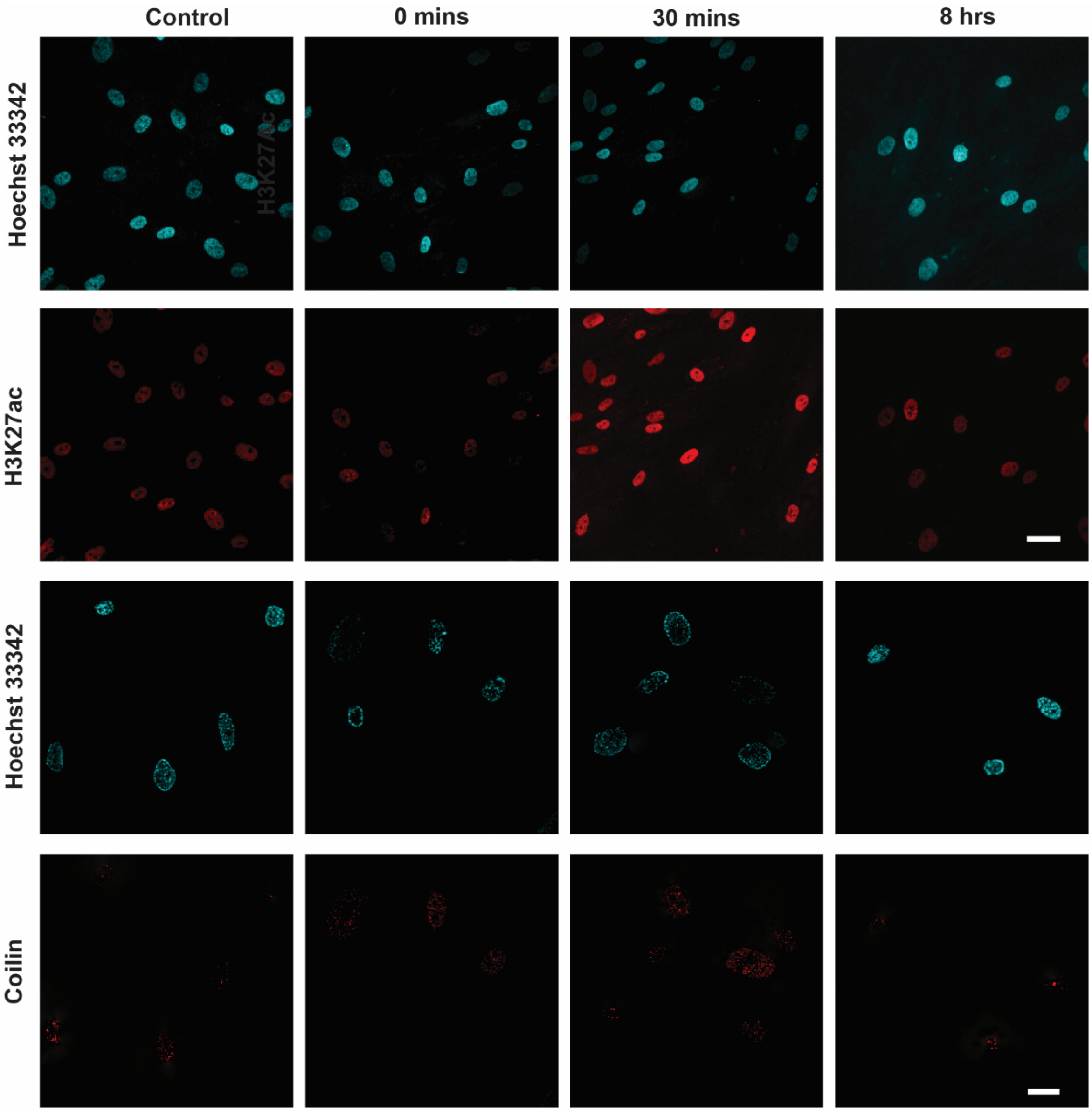


**Figure S7** Representative confocal microscopy images of the nucleus (stained with Hoechst 33342 and displayed in cyan) together with H3K27ac3 (displayed in red) and Coilin (displayed in red) immunofluorescence following SRBW nanomechanostimulation of the hMSCs at different post-exposure incubation times, compared to those of the control (unstimulated) cells. The scale bars denote lengths of 40 µm.

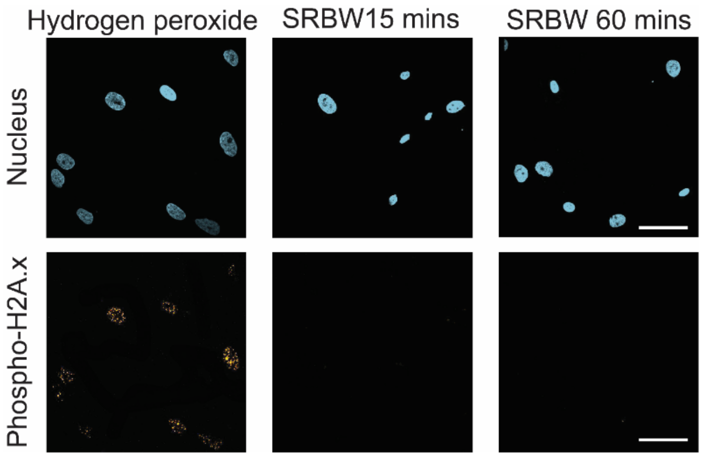


**Figure S8** Representative confocal microscopy images (60x magnification) of hMSCs stained for their nuclei with Hoechst 33342 (blue) and with phosphor-H2A.X—a marker for DNA damage (displayed in yellow), when treated with either hydrogen peroxide ($H_2O_2$) or the SRBW nanomechanostimulation. In the former, the cells were incubated with $H_2O_2$ for 20 mins, followed by incubation with fresh media for a further 30 mins, whereas the SRBWnanomechanostimulated cells were fixed after 15 and 60 mins post-exposure incubation. The scale bars denote 50 µm lengths.

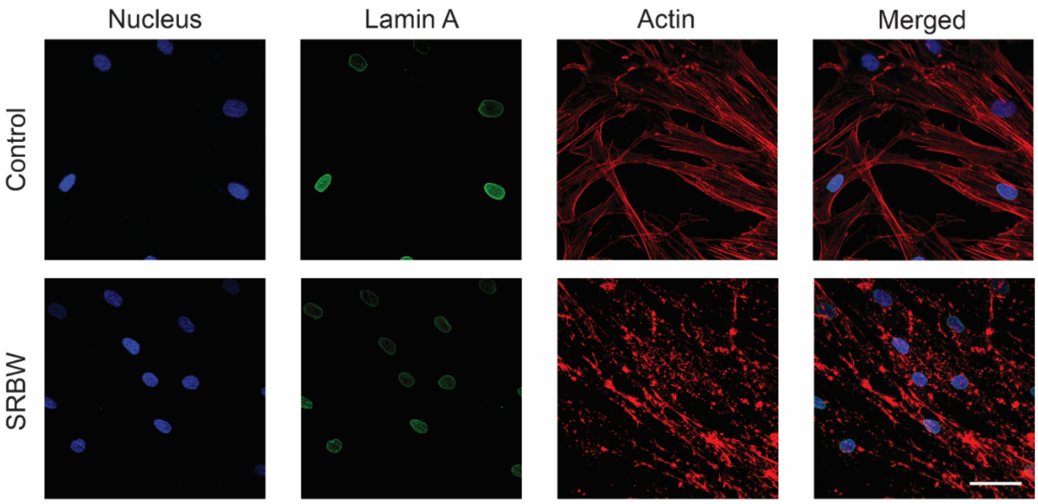


**Figure S9** Representative confocal microscopy images showing the effect of cytochalasin D on actin (displayed in red), Lamin A (displayed in green) and the nucleus (displayed in blue). The control (unstimulated) cells were incubated in DMSO (vehicle for cytochalasin D). The scale bar denotes a length of 50 µm.

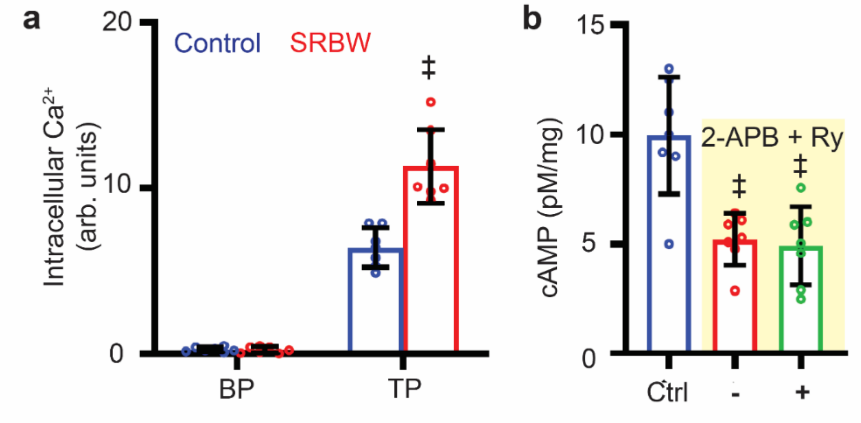


**Figure S10** (a) Intracellular $Ca^{2+}$ concentration ($n$ = 7) in the presence of the intracellular calcium chelator BAPTA-AM (BP) and the SERCA inhibitor thapsigargin (TP) in control (unstimulated) cells (blue) and cells subjected to the SRBW nanomechanostimulation (red), with respect to cells without BAPTA-AM treatment. (b) Intracellular cAMP concentration ($n$ = 7) in control (unstimulated) cells (blue), and SRBW nanomechanostimulated cells, both in the absence (–; red) and in the presence (+; green) of ER calcium efflux pump (IP3R and RyR) inhibitors 2-APB and Ry. The data are represented in terms of the mean value ± the standard error over triplicate runs. ‡ indicates statistically significant differences with $p$ < 0.0001.

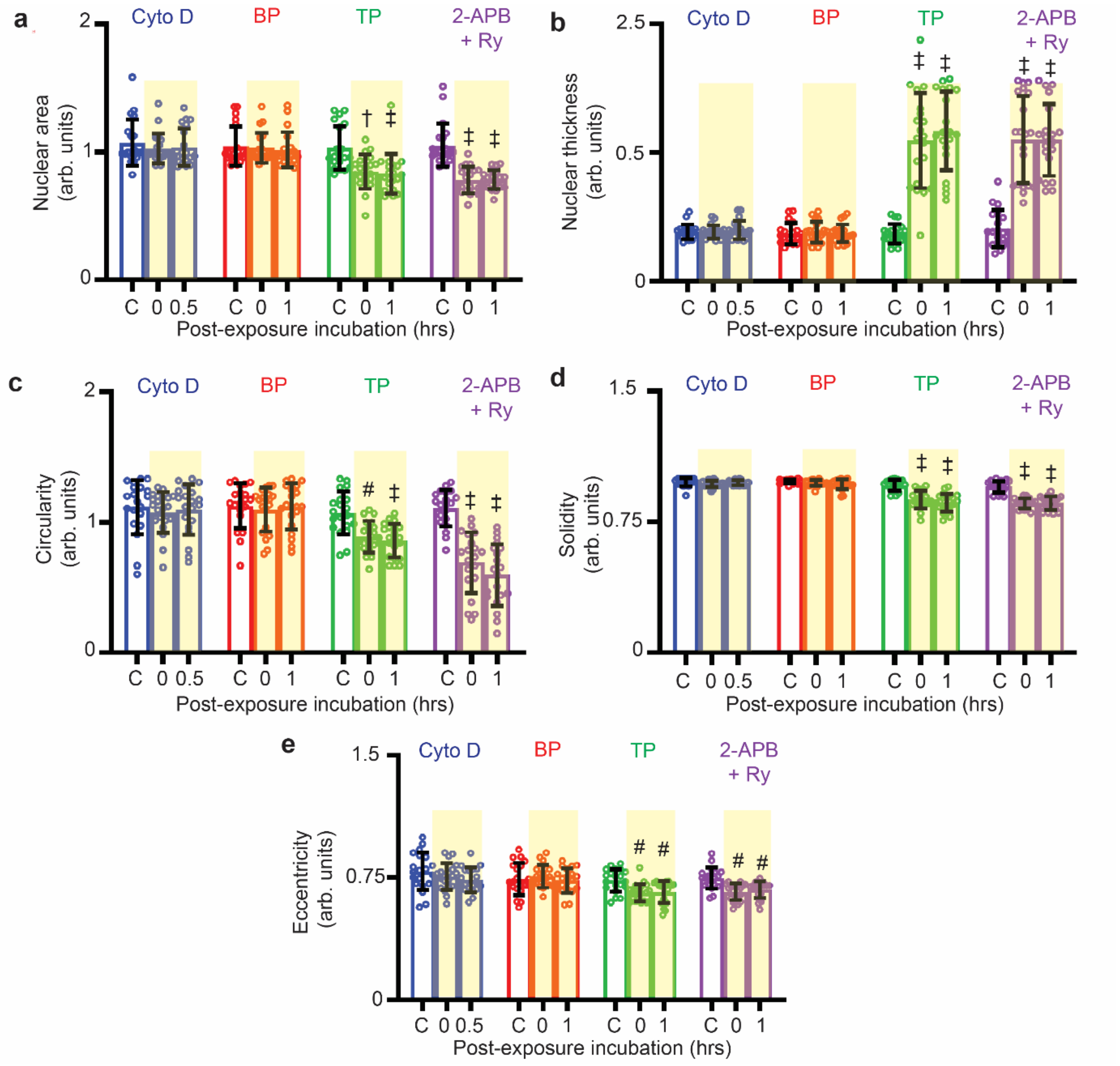


**Figure S11** Quantification of nuclear morphometry (i.e., nuclear (a) area, (b) thickness, (c) circularity, (d) solidity, and, (e) eccentricity)) changes from inhibitor/chelator studies at 0 and 30 mins or 1 hr post-exposure incubation following SRBW nanomechanostimulation, compared to that in the control (unstimulated) cells (C). Data are represented in terms of the mean value ± the standard error over multiple runs (*n* = 20 from four independent experiments); #, † and ‡ indicate statistically significant differences with $p < 0.01$, $p < 0.001$, and $p < 0.0001$, respectively.

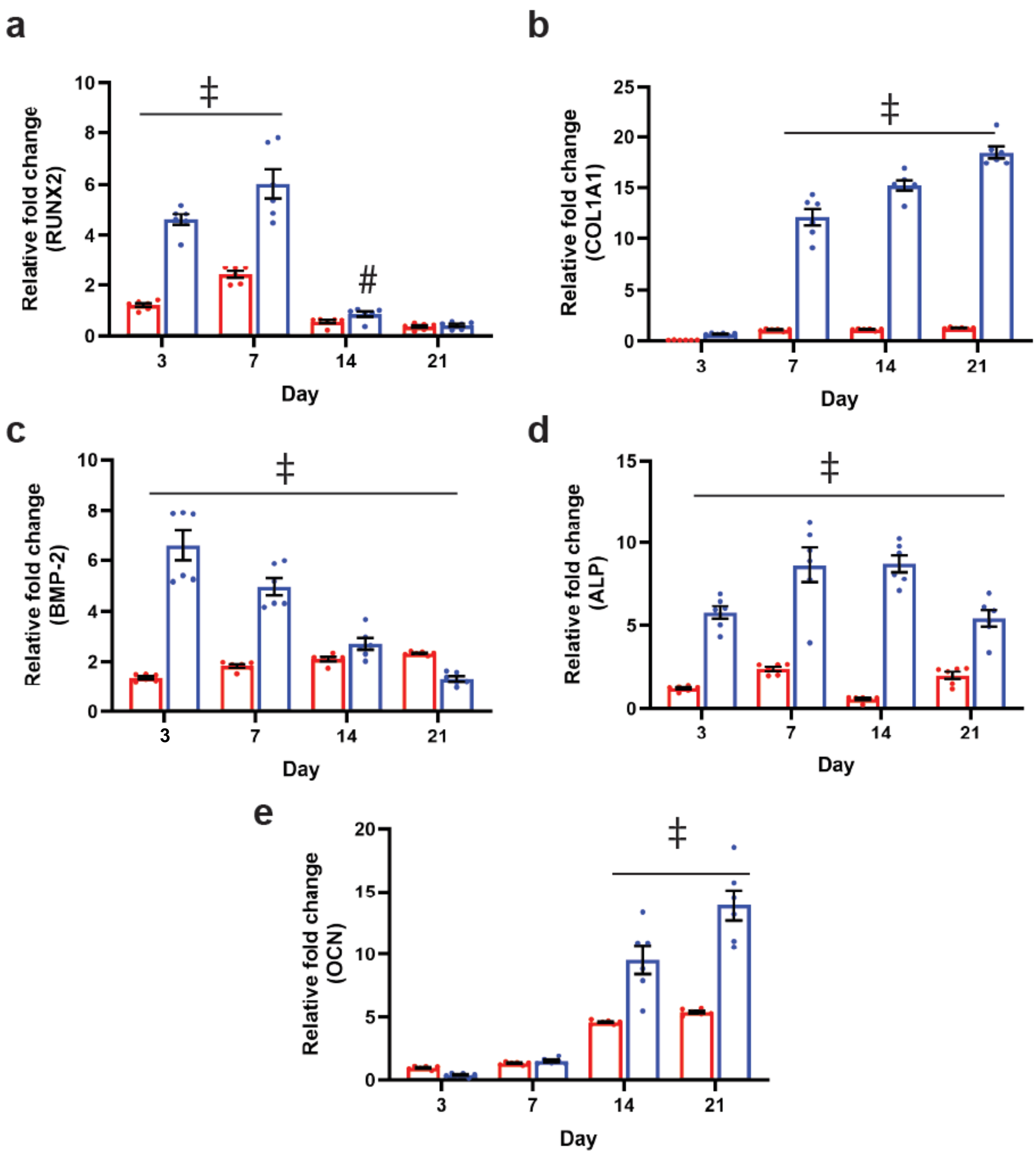


**Figure S12** mRNA profiling of (a) RUNX2, (b) COL1A1, (c) BMP-2, (d) ALP, and, (e) OCN, showing the relative fold change in gene expression (normalized against that of unstimulated cells in osteogenic media (OM); positive control) at different time points obtained for unstimulated hMSCs in osteogenic media (red), and, SRBW-nanomechanostimulated hMSCs in basal media (blue). The data are represented in terms of the mean value ± the standard error; ‡ indicates statistically significant differences with $p < 0.0001$.

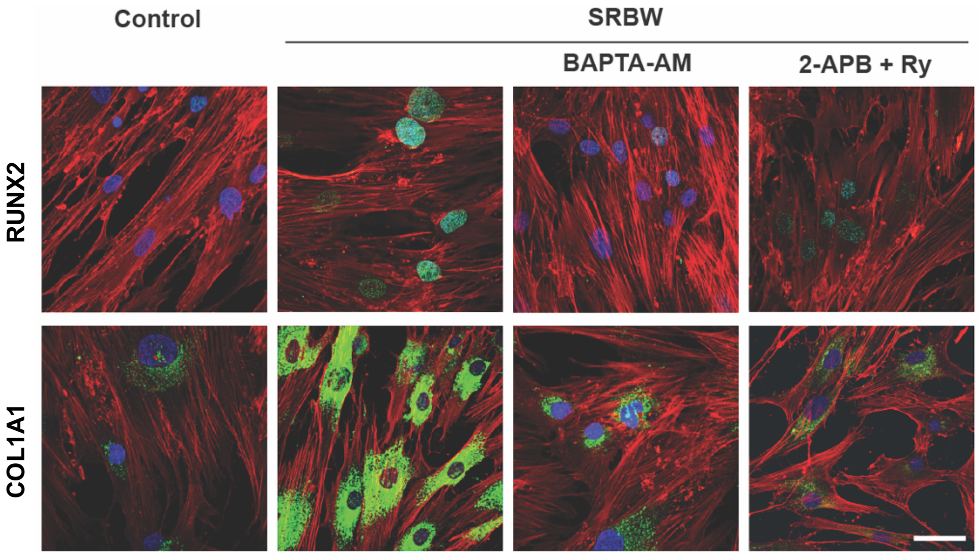


**Figure S13** Representative confocal microscopy images (day 3) showing RUNX2 and COL1A1 expression (displayed in green) in unstimulated hMSCs in basal media (negative control), and SRBW nanomechanostimulated hMSCs in basal media, in the absence and in the presence of an intracellular $Ca^{2+}$ chelator (BAPTA-AM) and an IP3R and RyR inhibitor (2-APB + Ry). Nuclei were stained with Hoechst 33342 and are displayed in blue, whereas actin was stained using phalloidin (ActinRed$^{TM}$ 555) and displayed in red. The scale bar denotes a length of 100 µm.

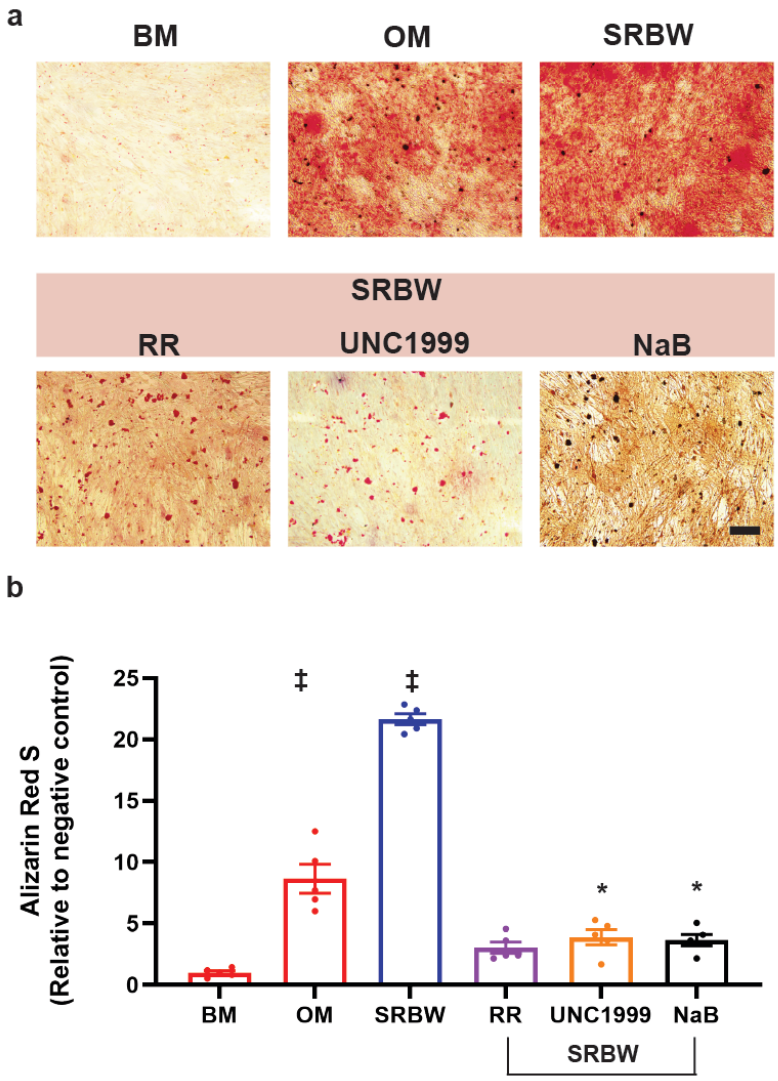


**Figure S14** (a) Left to right: Representative light microscopy images showing alizarin red staining (day 17) of unstimulated hMSCs in basal (negative control; BM) and osteogenic (positive control; OM) media, and, hMSCs treated with the SRBW nanomechanostimulation in basal media in the absence and in the presence of various inhibitors: a Piezo channel inhibitor ruthenium red (RR), a histone methylation inhibitor UNC1999, and a histone deacetylase inhibitor sodium butyrate (NaB). The scale bar denotes a length of 20 µm. (b) Extent of mineralization (extracted using 10% acetic acid) at day 17 of the aforementioned cases, normalized against that of the negative control (BM); data are represented in terms of the mean value ± the standard error; * and ‡ indicate statistically significant differences with $p < 0.05$ and $p < 0.0001$, respectively.

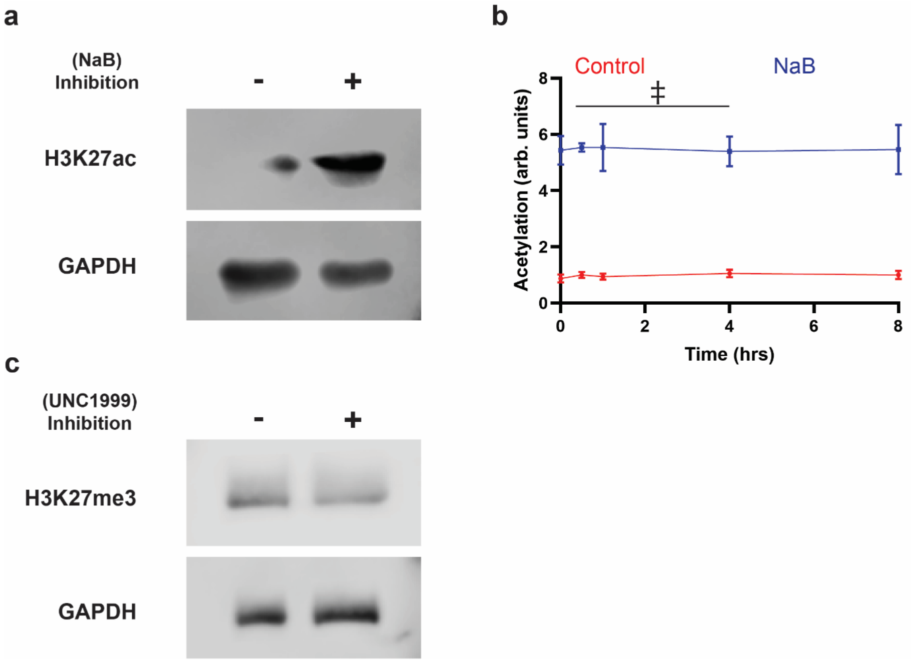


**Figure S15** (a) Western blots of (a) H3K27ac and (c) H3K27me3 for uninhibited and inhibited (with a histone methylation inhibitor UNC1999, and a histone deacetylase inhibitor sodium butyrate (NaB)) hMSCs in basal media along with that of the loading control (GAPDH). (b) Quantification of H3 acetylation in hMSCs by ELISA with time for the case when the cells are inhibited with NaB (normalized against the case of unstimulated cells in basal media, i.e., the negative control); data are represented in terms of the mean value ± the standard error; ‡ indicates statistically significant differences with $p < 0.0001$.

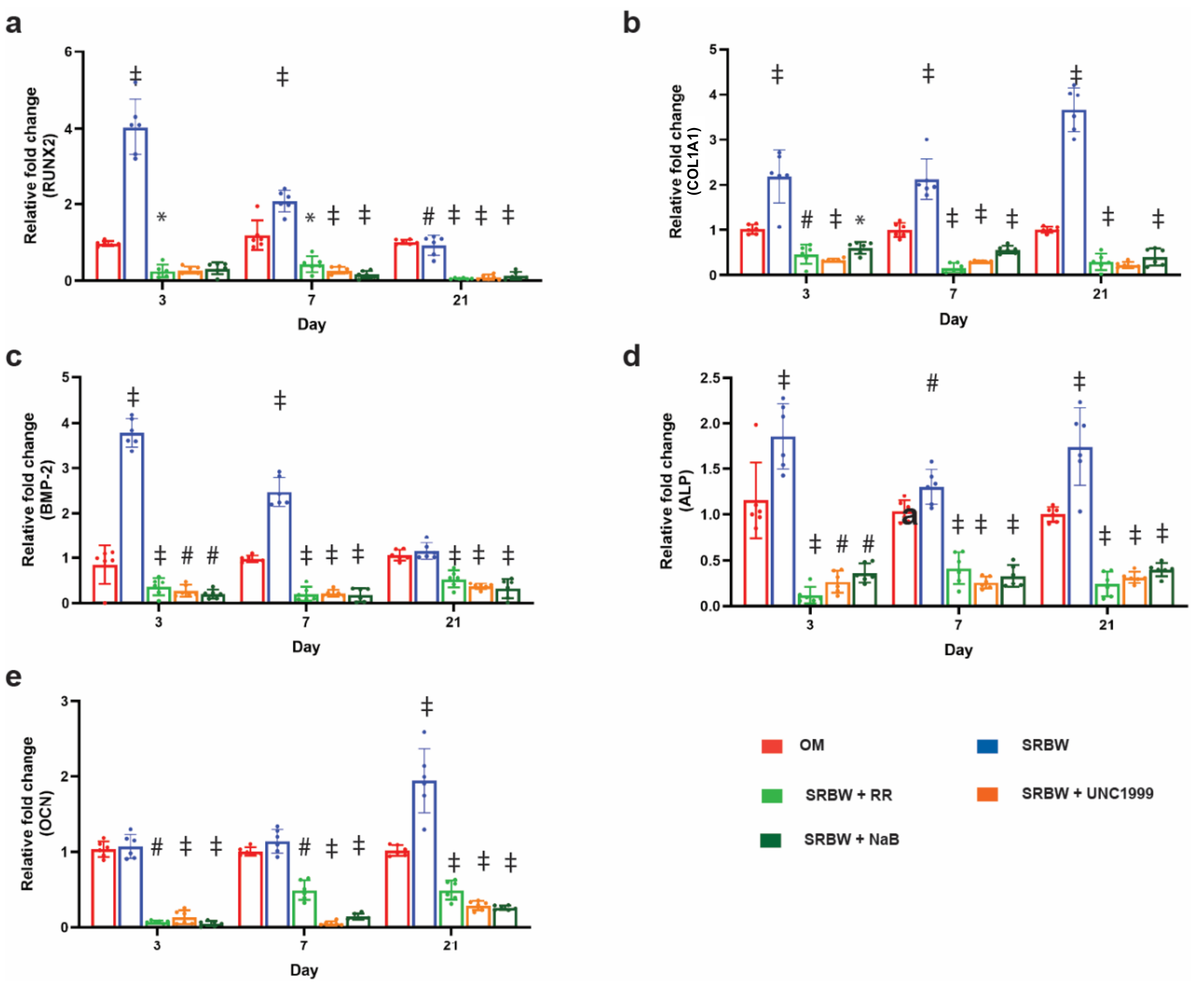


**Figure S16** mRNA qPCR of (a) RUNX2, (b) COL1A1, (c) BMP-2, (d) ALP, and, (e) OCN, showing the relative fold change in gene expression (against that of unstimulated cells in osteogenic media (OM); positive control) at different time points obtained for unstimulated hMSCs in osteogenic media, and, SRBW-nanomechanostimulated hMSCs in basal media in the absence and in the presence of various inhibitors: a Piezo channel inhibitor ruthenium red (RR), a histone deacetylase inhibitor sodium butyrate (NaB) and a histone methylation inhibitor UNC1999. Cells were incubated with the inhibitors for 5 days as with the SRBW nanomechanostimulation. The data are represented in terms of the mean value ± the standard error; # and ‡ indicate statistically significant differences with $p < 0.01$ and $p < 0.0001$, respectively.

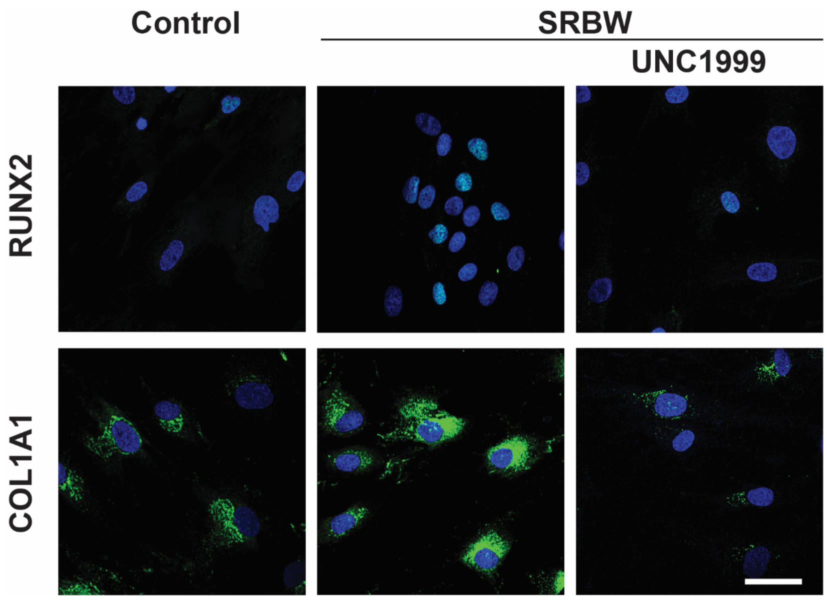


**Figure S17** Representative confocal microscopy images (day 3) showing RUNX2 and COL1A1 expression (displayed in green) in unstimulated hMSCs in basal media (negative control), and SRBW nanomechanostimulated hMSCs in basal media, in the absence and in the presence of a histone methylation inhibitor UNC1999. Nuclei were stained with Hoechst 33342 and displayed in blue. The scale bar denotes a length of 100 µm.
